# Ectopic HCN4 expression drives mTOR-dependent epilepsy

**DOI:** 10.1101/853820

**Authors:** Lawrence S. Hsieh, John H. Wen, Lena H. Nguyen, Longbo Zhang, Juan Torres-Reveron, Dennis D. Spencer, Angélique Bordey

## Abstract

The causative link between focal cortical malformations (FCM) and epilepsy is well-accepted, especially among patients with focal cortical dysplasia type II (FCDII) and tuberous sclerosis complex (TSC). However, the mechanisms underlying seizures remain unclear. Using a mouse model of TSC- and FCDII-associated FCM, we show that FCM neurons are responsible for seizure activity via their unexpected abnormal expression of the hyperpolarization-activated cyclic nucleotide-gated potassium channel isoform 4 (HCN4), which is normally not present in cortical pyramidal neurons after birth. Increasing intracellular cAMP levels, which preferentially affects HCN4 gating relative to the other isoforms, drove repetitive firing of FCM neurons but not that of control pyramidal neurons. Ectopic HCN4 expression was mTOR-dependent, preceded the onset of seizures, and was also found in diseased neurons in tissue resected for epilepsy treatment from TSC and FCDII patients. Finally, blocking HCN4 channel activity in FCM neurons prevented epilepsy in mice. These findings that demonstrate HCN4 acquisition as seizure-genic, identify a novel cAMP-dependent seizure mechanism in TSC and FCDII. Furthermore, the unique expression of HCN4 exclusively in FCM neurons provides opportunities for using HCN4 as a gene therapy target to treat epilepsy in individuals with FCDII or TSC.

**One Sentence Summary:** Our data provide a novel cAMP-dependent mechanism of seizure initiation in focal cortical dysplasia and tuberous sclerosis complex due to the unexpected ectopic expression of HCN4 channels only in diseased neurons. HCN4 channels are thus promising candidates for gene therapy to treat epilepsy generated by mTOR-driven focal malformations.

## Introduction

Disorders caused by mutations in the mTOR pathway genes lead to mTOR hyperactivity, brain malformations, and life-long epilepsy in the majority of patients (1–3). Among this group of disorders, FCD and TSC account for the largest population of children and young adults that undergo brain surgery to treat intractable epilepsy (4–6). Nearly two-thirds of these patients are refractory to treatment with antiepileptic drugs and experience life-long seizures, leading to a spectrum of neurocognitive and psychological disabilities (1). Upon failure of conventional anti-epileptic drugs, surgical resection of the seizure foci or pharmacological inhibition of mTOR signaling in the case of TSC are the only available treatments for mTOR-dependent focal epilepsy, but neither option is fully effective (7–9). Despite the identification of the molecular pathways dysregulated in TSC and FCDII and the pathological characteristics associated with seizures (4, 9–11), the mechanism of epileptogenesis remains unknown, although several hypotheses of epileptogenicity have been proposed. These include abnormal reorganization of the circuitry within and surrounding FCMs and abnormal electrophysiological properties of FCM neurons (5, 12–15). However, none of these mechanisms has been tested. In addition, the exact site and the cell types responsible for epileptogenic activity in humans remain controversial as some studies have recorded epileptic discharges both inside and outside of FCMs whereas other studies have found FCMs to be electrically silent (14, 15). The lack of consensus on the site of seizure generation further reflects our poor understanding of the epileptogenic mechanisms, which has hampered the identification of novel therapeutic targets and effective treatment options.

Using a mouse model of TSC and FCDII (16), we show that silencing FCM neurons prevents the occurrence of seizures, suggesting that FCM neurons are responsible for seizure activity. To identify how FCM neurons drive seizures, we obtained patch clamp recordings in FCM neurons and unexpectedly identified the abnormal expression of HCN4, which is not present in normal pyramidal neurons. While FCM neurons were depolarized and hence closer to the threshold for action potential generation, they required significantly more current injection than in controls to reach that threshold due to the cells’ increased size and intrinsic conductance. Consistent with the fact that HCN4 channels are the most cAMP sensitive of all HCN channels (17–19), increasing intracellular cAMP induced repetitive firing of FCM neurons and provides a novel mechanism of firing and seizure induction compared to the traditional excitatory input-induced firing mechanism. Importantly, HCN4 was also found in dysmorphic cortical neurons from TSC and FCDII patients who underwent surgery for epilepsy treatment. Finally, using a genetic approach to block HCN4 activity in FCM neurons prevented seizures. Collectively, our findings identify a novel mechanism of seizures in mTOR-dependent FCM that provide the first gene therapy treatment option to prevent seizure initiation in patients with TSC and FCDII.

## Results

### Silencing FCM neurons decreases seizure activity

To investigate the mechanisms of seizure generation in FCM, we used a mouse model that recapitulates the focal nature of the cortical malformation and the cellular mosaicism of human FCDII and TSC brains (16, 20). To generate this model, we used *in utero* electroporation (IUE) to induce mTOR hyperactivity in neurons in a limited cortical area at embryonic day (E) 15 (Fig. 1A) (16, 20–24). To increase mTOR activity, we expressed a constitutively active mutant Rheb (Rheb^CA^), the canonical mTOR activator (21, 23, 25–27) (Fig. 1B). A fluorescent protein (GFP or tdTomato) was co-expressed to label neurons. These mice developed a singular FCM in the medial prefrontal cortex (Fig. 1C) that displayed a pathology resembling that of human FCM, including the presence of misplaced and cytomegalic Rheb^CA^-expressing neurons that are interchangeably called FCM neurons (Fig. 1D and E). These FCM neurons showed a significant increase in soma size and immunoreactivity for phosphorylated protein S6 (phospho-S6), a read-out of mTOR activity, compared to control neurons in mice electroporated with GFP only (Fig. 1F and G). Mice containing FCM were visually observed to develop convulsive seizures by postnatal day (P) 21. To better characterize the seizures, we obtained epidural electroencephalography (EEG) recordings combined with video monitoring for 5-7 consecutive days in >2 month-old mice. Mice containing FCM exhibited recurrent, Racine grade 4-5 seizures with classical interictal, tonic, clonic, and postictal periods (Fig. 1H and I, and movie S1).

**Figure 1:**
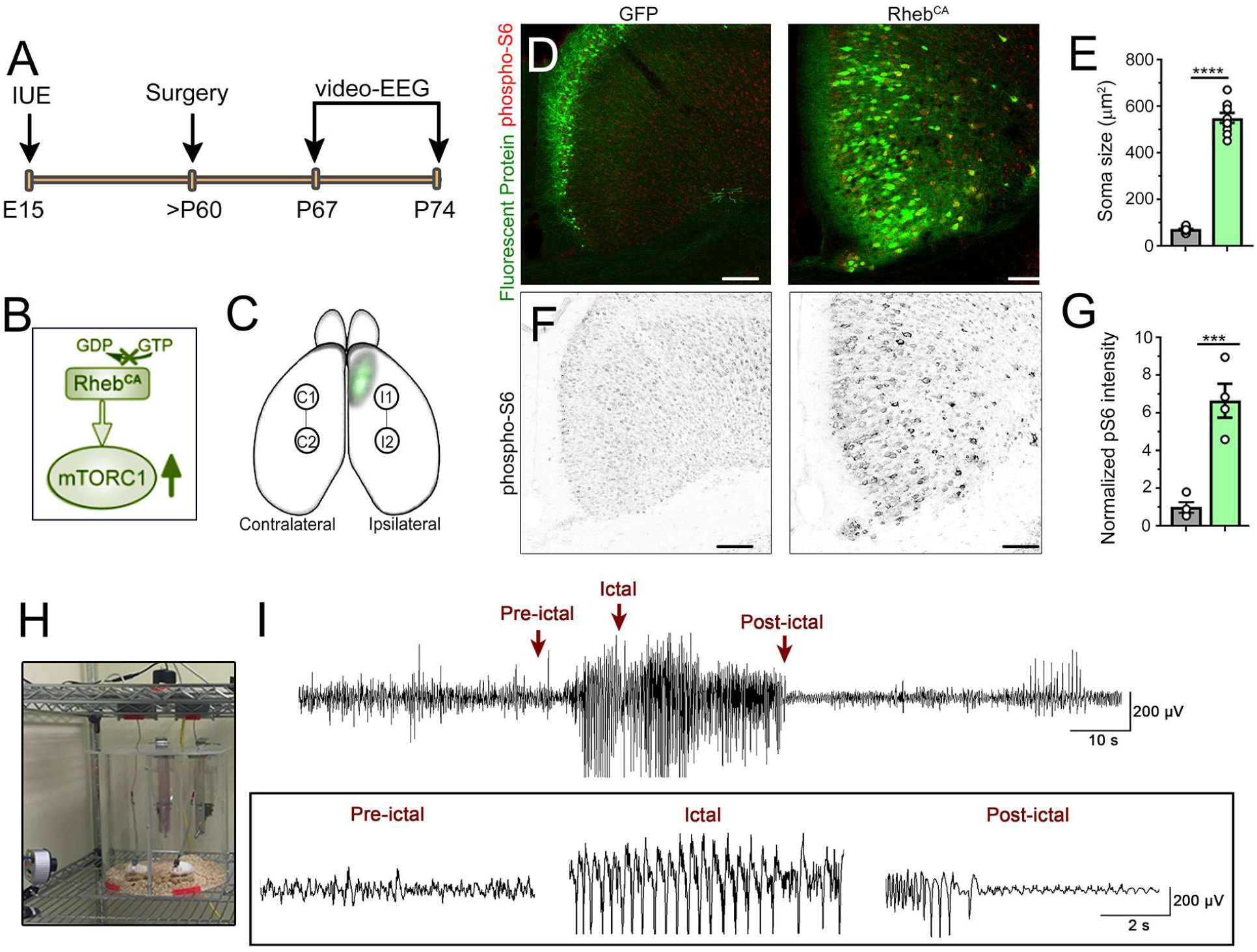
Model of mTOR-dependent FCM-associated seizures. **(A)** Timeline of the experimental paradigm. **(B)** Diagram of Rheb^CA^ effect on mTOR complex 1 (mTORC1). **(C)** Diagram of a mouse brain with superimposed image of a FCM in the medial prefrontal cortex (mPFC). Linked circles mark the approximate locations of independent pairs of recording electrodes in the ipsilateral and contralateral hemispheres. **(D)** Confocal images of GFP and tdTomato (pseudo-colored green) (noted fluorescent proteins) fluorescence and phospho-S6 immunostaining (red) in coronal sections from 3 months old mice expressing GFP (control) or Rheb^CA^ (+ tdTomato) in the mPFC. Scale bars: 150 µm. **(E)** Bar graphs of soma size of GFP and Rheb^CA^-expressing neurons. Student’s t test. **(F)** B&W phospho-S6 immunostaining from images shown in D. Scale bar: 150 µm. **(G)** Bar graphs of phospho-S6 (pS6) immunofluorescence in GFP and Rheb^CA^-expressing neurons. Student’s t test. **(H)** Image of the video-EEG set-up. **(I)** Representative examples of an EEG trace and higher temporal resolution traces in the inset.

Using our mouse model of FCM, we previously reported that FCM neurons display depolarized resting membrane potentials (RMP) compared to their control counterparts and are therefore closer to the threshold for generating action potentials (21). We thus examined whether silencing FCM neurons without normalizing their morphological abnormalities, which could contribute to seizure generation, would limit the incidence of seizures. To do this, we overexpressed inwardly rectifying potassium channels Kir2.1 that are expected to hyperpolarize neurons expressing these channels as well as decrease their membrane resistances (28). Using E15 IUE, we co-expressed Kir2.1 with Rheb^CA^ (Fig. 2A). In the control condition, GFP was co-electroporated with Rheb^CA^. Kir2.1 expression in Rheb^CA^ neurons did not prevent the formation of FCM (Fig. 2A and B) and did not interfere with mTOR hyperactivity as shown by quantification of soma size and phospho-S6 immunofluorescence, respectively (Fig. 2C). Furthermore, patch clamp recordings of Rheb^CA^ neurons in acute slices from P21 to P35 validated that Kir2.1 expression did not alter membrane capacitance (a read-out of soma size), but it significantly hyperpolarized Rheb^CA^ neurons and increased their membrane conductance (Fig. 2D). The properties of current-induced action potentials, including half-width and threshold, were unaltered by Kir2.1 expression (Fig. 2E and F), but consistent with the changes in RMP and membrane conductance, larger current injections were required for the generation of action potentials Kir2.1-expressing FCM neurons (Fig. 2G). This resulted in a significant shift in the input (injected current amplitude)-output (number of action potentials) curve (Fig. 2H). Thus, expressing Kir2.1 renders FCM neurons less excitable, in part via hyperpolarizing their RMP further away from the action potential threshold. Importantly, mice overexpressing Kir2.1 in FCM neurons displayed a significant decrease in seizure frequency (from a mean of 4.5 to 0.5 seizures per day) without affecting seizure duration (Fig. 2I and J). Collectively, these data show that alterations in the electrical properties of FCM neurons are necessary for seizure initiation and that silencing these neurons is sufficient to alleviate seizures.

**Figure 2:**
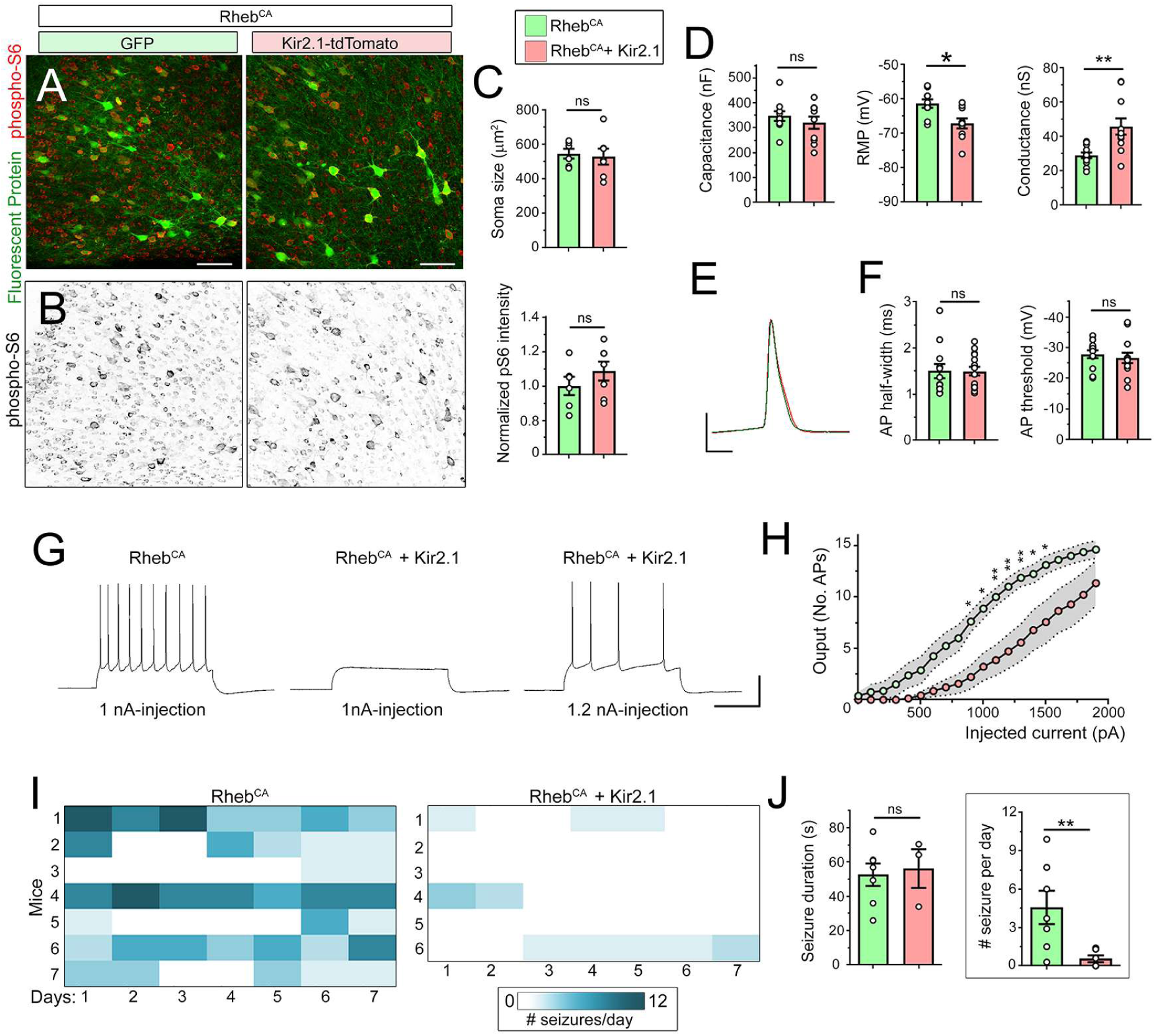
Silencing cytomegalic neurons in mTOR-driven FCM prevents seizure activity. **(A)** Confocal images of GFP and tdTomato fluorescence (pseudo-colored green) and phospho-S6 immunostaining (red) in coronal sections from 4 months old littermate mice (used in panel J) expressing Rheb^CA^ + GFP or + Kir2.1 (fused to tdTomato). Scale bar: 100 µm. **(B)** B&W phospho-S6 immunostaining from images shown in A. **(C)** Bar graphs of normalized phospho-S6 immunofluorescence intensity and soma size of Rheb^CA^ neurons co-expressing GFP (green) or Kir2.1 (pink). Student’s t test. **(D)** Bar graphs of cell capacitance, resting membrane potential (RMP), and membrane conductance of Rheb^CA^ neurons co-expressing GFP or Kir2.1. Patch clamp recordings were obtained in acute slices from P26-P42 mice. Student’s t test. **(E)** Superimposed individual action potentials from Rheb^CA^ neurons in both conditions. Scale bars: 2 ms/40 mV. **(F)** Bar graphs of the action potential (AP) half-width and threshold. Student’s t test. **(G)** Representative depolarization and action potentials upon current injection in Rheb^CA^ co-expressing GFP or Kir2.1. Scales: 200 ms/40 mV. **(H)** Injected current amplitude plotted as a function of the mean number of action potentials for generating an input-output curve in Rheb^CA^ neurons. Two-way repeated measure ANOVA, followed by Sidak post-test. The grey area outlines the SEM for each curve. **(I)** Heat map of the number of seizures over a 7-day long recording period in mice containing Rheb^CA^ neurons with GFP or Kir2.1. **(J)** Bar graphs of the duration and number of seizures per day in the two conditions. Student’s t test (seizure frequency) and Mann Whitney U test (seizure duration). Data are mean ± SEM. *: P<0.05, **: P<0.01, ***: P<0.001, ****: P<0.0001, ns: not significant. Exact *P* values can be found in Supplementary Table 4.

### Aberrant HCN channel expression and increases in cAMP depolarizes FCM neurons

We next examined the electrical properties of FCM neurons using patch clamp recordings in acute brain slices from P26-P42 mice. As we previously reported, we confirmed that FCM neurons displayed significantly depolarized RMPs and increased conductance compared to control neurons in littermate mice expressing only GFP (Fig. 3A). Consistent with an increased conductance, there was a significant shift in the input (injected current from their RMP)-output (# of action potentials) curve, suggesting that Rheb^CA^ neurons are less likely to generate action potentials upon depolarizations (Fig. 3C and D). Using hyperpolarizing current step, one unexpected observation was the present of a robust “sag” response in Rheb^CA^ neurons that was either minimal or mostly not present in control neurons (Fig. 3E) (29). Such a sag response suggests the presence of HCN channels. Under control conditions, HCN channels are primarily expressed in deep layer neurons but not in superficial layer (2/3) neurons (29), which are the neuronal population targeted by IUE at E15. In addition, HCN currents are known to control neuron RMPs and may thus contribute to the depolarized RMP in Rheb^CA^ neurons (17–19, 30). We thus preferentially focused on these channels. To validate that these sags were due to the presence of HCN channels, we recorded Rheb^CA^ neurons in the presence of the HCN channel blocker, zatebradine. Zatebradine eliminated the sag response in Rheb^CA^ neurons (Fig. 3F). Using voltage-clamp, we applied 500 ms-long hyperpolarizing voltage-steps (from −40 to −130 mV) to activate HCN channels (Fig. 3G). Rheb^CA^ neurons displayed large hyperpolarization-activated inward currents that were much larger than in the control neurons (Fig. 3G and H). These inward currents displayed a slow activation kinetics resembling that of HCN currents requiring a 3 s-long voltage pulse to reach full amplitude (Supplementary Fig. S1) and were significantly reduced in the presence of zatebradine (Fig. 3G and H). The increased hyperpolarization-induced inward currents were nevertheless not fully blocked by zatebradine presumably due to an increase in the amplitude of inwardly rectifying K currents (Kir) in Rheb^CA^ neurons compared to control due to the cell size increase (Fig. 3G and H). As evident on the current-membrane potential curves (Fig. 3H) and further quantified, zatebradine shifted RMP of cells to hyperpolarized values similar to those of control neurons (Fig. 3I). To quantify the h current amplitudes, we measured the amplitude difference (ΔI) between the onset and the end of the current trace using a 500 ms-long voltage step (see dotted line and Ih in Fig. 3G). Zatebradine significantly reduced these ΔI and there was no residual inward current at −90 mV (Fig. 3J). Throughout the rest of the manuscript, Ih current amplitude will then be measured as ΔI at −90 mV using this method. Consistent with the normalization of RMP with zatebradine, larger h currents were associated with more depolarized resting membrane potentials in Rheb^CA^ neurons (Fig. 3K). The amplitude of the h currents was also dependent on the amount of electroporated Rheb^CA^ plasmid and thus mTOR activity level (Supplementary Fig. S1). HCN channels contribute to the generation of rhythmic firing in neurons and heart cells (31–34) and they display different sensitivity to intracellular cAMP levels (17–19). We thus examined whether increasing cAMP in Rheb^CA^ neurons would be sufficient to trigger spontaneous, repetitive firing independently of depolarizations. We bath applied a well-characterized cell-permeable adenylate cyclase activator, forskolin, to increase in intracellular cAMP in acute brain slices containing Rheb^CA^ neurons. When recorded at the resting membrane potential, all Rheb^CA^ neurons were depolarized by forskolin and 4/9 neurons generated repetitive action potentials (Fig. 3L and M). This effect did not occur in control superficial layer neurons (Fig. 3L and M), which express very few or no HCN channels. Collectively, these data indicate that Rheb^CA^ neurons express ectopic HCN channels and the abnormal presence of these channels is responsible for the depolarized resting membrane potentials of Rheb^CA^ neurons and their repetitive firing upon increases in intracellular cAMP.

**Figure 3:**
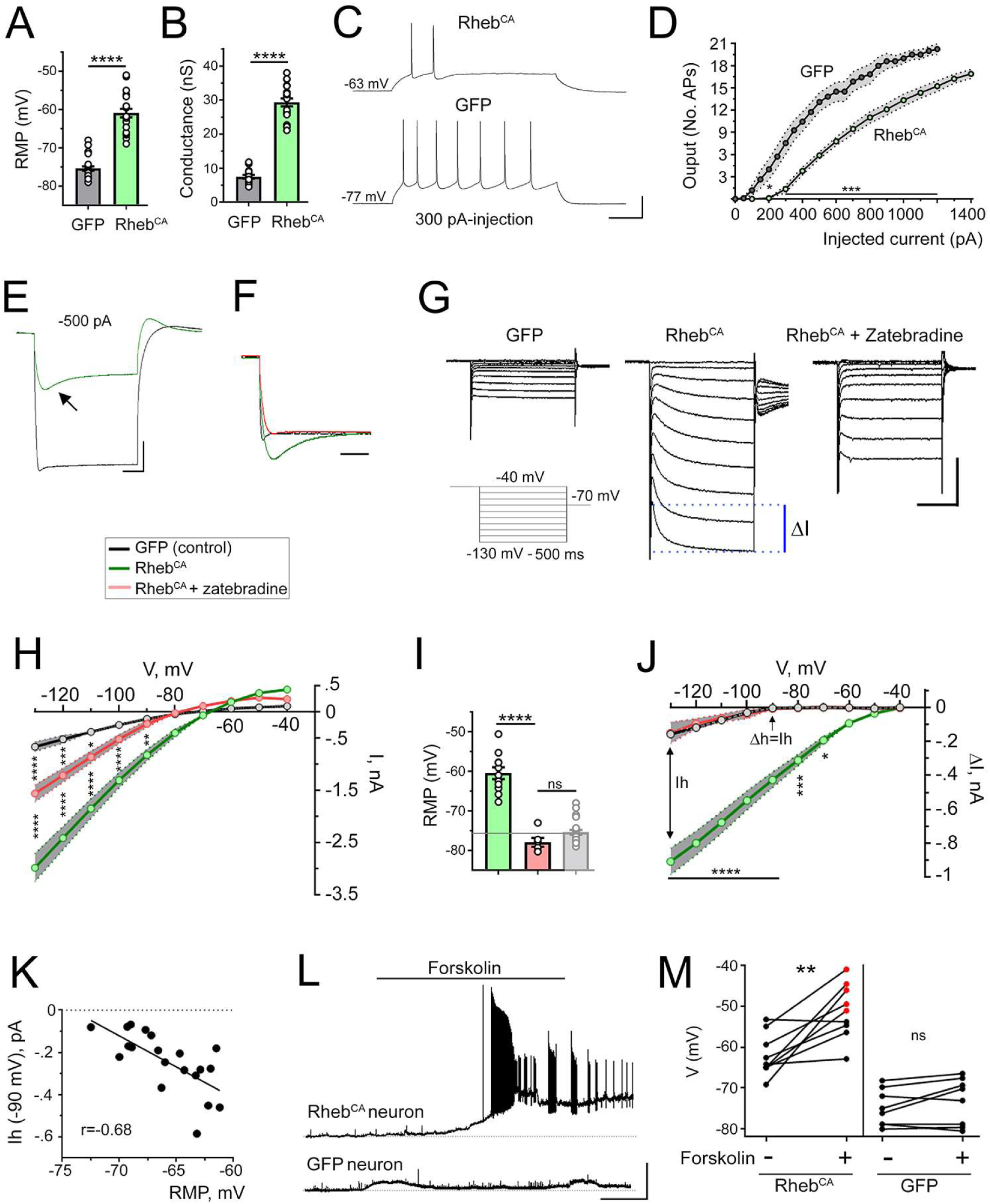
Abnormal HCN currents in FCM neurons are depolarizing and confer cAMP-dependent firing. **(A and B)** Bar graphs of the RMP and conductance of control (GFP) and Rheb^CA^ neurons recorded in littermate P26-P42 mice. Student’s t test. **(C)** Representative depolarization and action potentials upon current injection in Rheb^CA^ or control (GFP) neurons. Scales: 100 ms/40 mV. **(D)** Injected current amplitude plotted as a function of the mean number of action potentials for generating an input-output curve in Rheb^CA^ neurons. Two-way repeated measure ANOVA, followed by Sidak post-test. The grey area outlines the SEM for each curve. **(E)** Representative voltage traces in response to a −500 pA current step in neurons expressing GFP (black) or Rheb^CA^ (green). Neurons were recorded in current-clamp at their RMP and voltage traces were superimposed post-recording. The arrow points to a hyperpolarization-induced voltage sag. Scales: 100 ms/10 mV. **(F)** Voltage traces in response to a −500 pA current step from GFP-expressing neurons (black), Rheb^CA^-expressing neurons (green), and Rheb^CA^-expressing neurons in the presence of zatebradine (40 µM, red). Voltage responses were rescaled and superimposed post-recording. Scale: 100 ms. **(G)** Representative current traces in cortical neurons expressing GFP (control), Rheb^CA^, or Rheb^CA^ with zatebradine (D). Protocol: conditioning step to −40 mV followed by 10 mV hyperpolarizing steps from −130 to −40 mV. Scale bars: 200 ms/1 nA. The blue dotted lines illustrate where the difference in current amplitude (ΔI) was measured within each voltage step to generate current-voltage (ΔI-V) curves in H. **(H)** Current amplitude (I measured at the end of the trace) versus the voltage in each condition. The grey area indicates the SEM for each curve. Two-way repeated measures ANOVA followed by Tukey post-test. **(I)** Bar graphs of the RMP for each condition. Control from panel A was added for comparison. Student’s t test. **(J)** ΔI-V curves. At −90 mV, ΔI corresponds to Ih. Two-way repeated measures ANOVA followed by Tukey post-test, Statistics is for Rheb^CA^ vs Rheb^CA^ + zatebradine. **(K)** Scatter plot of Ih (measured at −90 mV step) against the RMP. Two-tailed Pearson r with correlation coefficients. **(L and M)** Increasing intracellular cAMP via forskolin bath-application significantly depolarized Rheb^CA^ neurons but not control neurons, and it induced (triggered) repetitive (regenerative) firing in 4/9 Rheb^CA^ neurons. Scale bars: 5 min/30 mV. Paired Student’s t test. Data are mean ± SEM. ****:P<0.0001, ***:P<0.001, *:P<0.05, and ns: not significant. Exact *P* values can be found in Supplementary Table 4.

### Ectopic HCN4 expression in FCM neurons

HCN channels are encoded by four genes, *HCN1-4*, with different expression patterns throughout the brain (17). In the adult cortex, deep layer pyramidal neurons predominantly express HCN1 and low levels of HCN2 at the protein level (35). HCN3 and HCN4 display weak, diffuse expression in the cortex (35). Recently, HCN4 expression has been found in neuronal cell bodies scattered in the cortex that may be GABAergic neurons, but it is nearly absent in the cortex of young adult mice (35–37). Furthermore, in situ hybridization and immunoblotting detect low HCN3 and HCN4 mRNA levels in the adult cortex (38–40). HCN4 followed by HCN2 channels are the most sensitive to intracellular cAMP levels while HCN1 channels display low sensitivity (17–19). We thus immunostained for HCN1-4 in 2 months old mouse brain sections containing FCM. Consistent with previous studies (35), we identified intense HCN1 staining predominantly in apical dendrites of deep layer neurons and weak, diffuse HCN2 staining in the cortex (Fig. 4A and B). However, we found no changes in HCN1 and HCN2 staining pattern in the ipsilateral cortex containing FCM compared to the contralateral cortex as well as no staining in Rheb^CA^ neurons (Fig. 4E and F). We found no HCN3 staining in the cortex (data not shown). However, we identified strong HCN4 immunostaining in the cortex containing Rheb^CA^ neurons using two different antibodies against HCN4 (Fig. 4C and D). There was no expression of HCN4 in neurons of the contralateral hemisphere suggesting that aberrant HCN channel expression does not result from recurrent seizures. One of the antibodies (Alomone) was previously validated in vivo in conditional knockout mice (36). In addition, the specificity of both HCN4 antibodies was further confirmed by RNAi followed by immunoblotting in vitro (Supplementary Fig. S2) and exclusive immunoreactivity in neurons electroporated with an HCN4 overexpression plasmid and not in non-electroporated neurons in the vicinity (Supplementary Fig. S3). About 85% of Rheb^CA^ neurons displayed HCN4 immunoreactivity that decorated their soma, dendrites, and axons (Fig. 4C, G, and I). Finally, GFP-expressing cells in mice electroporated with GFP instead of Rheb^CA^ did not display HCN4 expression (Fig. 4H and I). As a positive control, HCN4 staining was observed in the cerebellum as expected (Supplementary Fig. S4). Thus, Rheb^CA^-expressing neurons display selective HCN4 expression that was absent in control pyramidal neurons.

**Figure 4:**
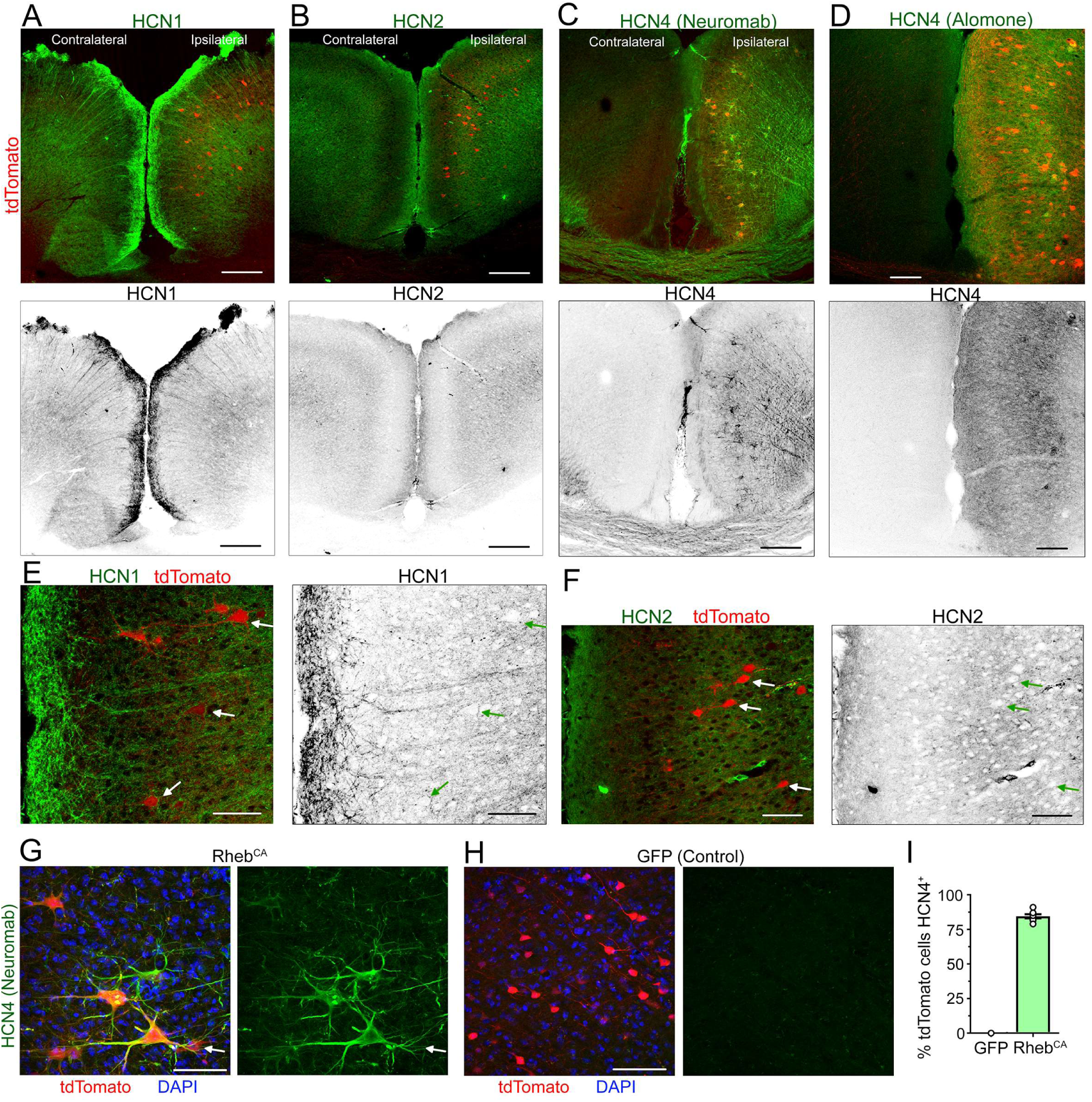
FCM neurons display abnormal mTOR-dependent HCN4 expression. **(A-D)** Immunostaining for HCN1 (A), HCN2 (B), and HCN4 (C) in coronal sections containing Rheb^CA^ neurons co-expressing tdTomato. HCN4 staining was performed using antibodies from Neuromab and Alamone. The bottom images display HCN staining in B&W. Scale bars: 200 µm. **(E and F)** Higher magnification images of HCN1 (E) and HCN2 (F) immunostaining (green and B&W) and tdTomato (red) from images in A and B. Scale bars: 60 µm. **(G)** Higher magnification images of HCN4 (green, Neuromab antibody), DAPI, and tdTomato. Scale bar: 60 µm. **(H)** Images of HCN4 immunostaining, tdTomato fluorescence and DAPI in GFP electroporated mice. Scale bar: 60 µm. **(I)** Bar graph of the percentage (%) of tdTomato neurons expressing HCN4 in mice electroporated at E15 with Rheb^CA^ or GFP.

### HCN4 expression is mTOR-dependent and precedes seizure onset

Higher magnification images illustrate that Rheb^CA^ neurons displaying HCN4 immunoreactivity also expressed increased phospho-S6 staining (Fig. 5A and B). To further assess whether increased mTOR activity was responsible for the abnormal expression of HCN4, we treated mice with the mTOR inhibitor rapamycin using the treatment paradigm (1 mg/kg every 48 hours from P1 to 2 months of age) that prevented FCM and the development of seizures (16). Rapamycin treatment prevented the expression of HCN4 in Rheb^CA^ neurons (Fig. 5C and D). Because HCN channel expression has been shown to be up- or down-regulated by seizures (for review see (41)), we examined whether Rheb^CA^ neurons would express HCN during postnatal cortical development prior to the onset of convulsive seizures that were visible in >P21 mice. Recordings in slices from P8-P12 mice showed that Rheb^CA^ neurons displayed zatebradine-sensitive HCN currents that were significantly greater than in control GFP neurons recorded in littermate mice (Fig. 5E-G). In addition, there was a significant increase in the HCN currents in Rheb^CA^ neurons during development from P6-12 to P28-P42 but no significant change in control neurons (Fig. 5H). These data indicate that mTOR hyperactivity drives aberrant expression of HCN4 in FCM neurons prior to the inset of convulsive seizures.

**Figure 5:**
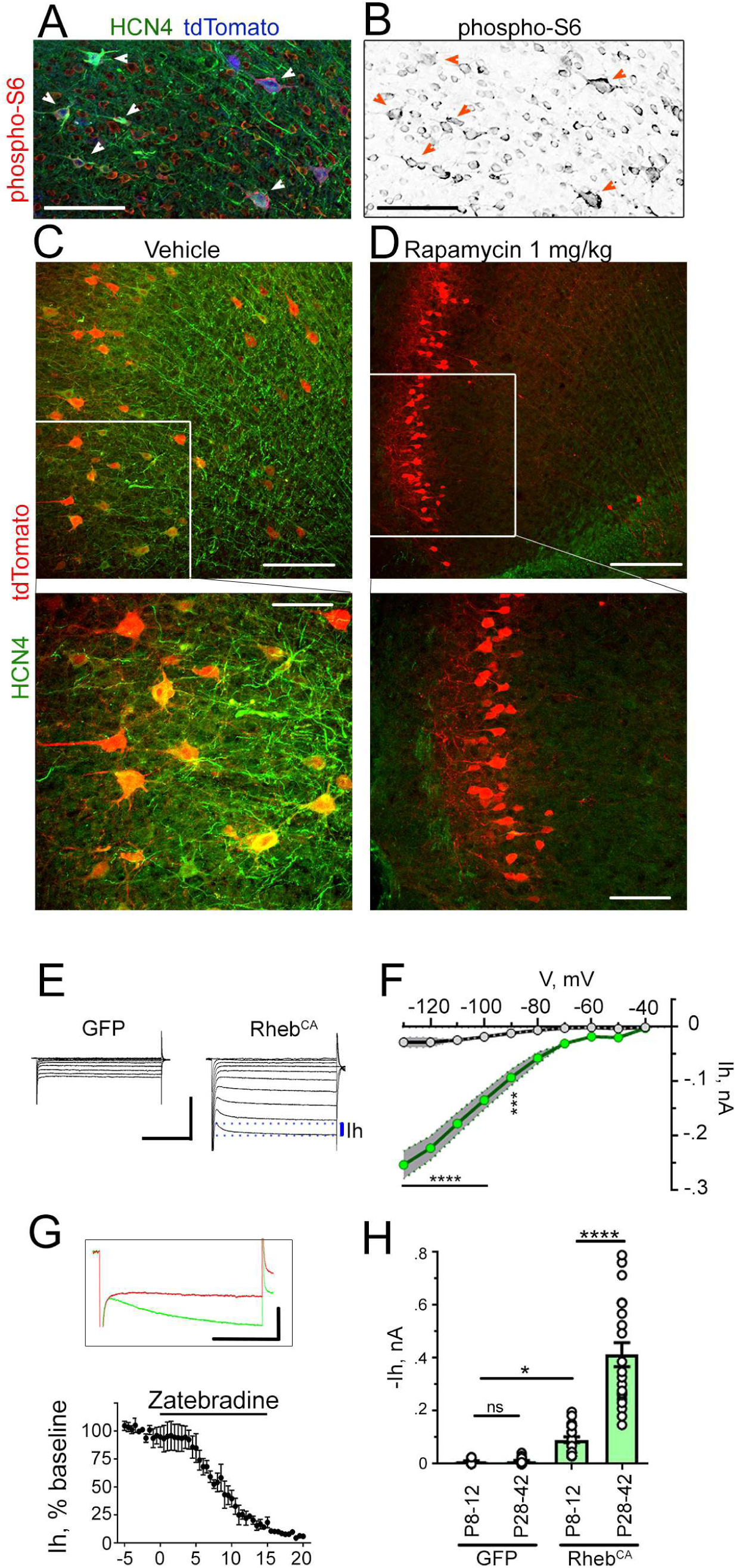
HCN4 expression is mTOR-dependent and precedes seizure onset. **(A and B)** HCN4 (Neuromab antibody) and phospho-S6 co-immunostaining in Rheb^CA^ neurons (arrowheads) co-expressing tdTomato (pseudo-colored blue). The bottom image illustrates phospho-S6 in B&W. Scale bars: 70 µm. **(C and D)** HCN4 immunostaining (Neuromab) and tdTomato fluorescence in Rheb^CA^ mice treated with vehicle (C) or rapamycin (D) and higher magnification images (from different number of optical sections from C). Scale bar: 30 µm. **(E and F)** Representative current traces in P8-P12 cortical neurons expressing GFP (control) or Rheb^CA^ (I). Scale bars: 200 ms/1 nA. The blue dotted lines illustrate where the h current amplitude (Ih) was measured within each voltage step to generate current-voltage (Ih-V) curves (J). Two-way repeated measures ANOVA followed by Tukey post-test. **(G)** Plot of the zatebradine block of Ih (measured at −90 mV) over time in a Rheb^CA^ neuron. Inset: Traces of Zatebradine block at −90 mV in Rheb^CA^ neurons. Scale bars: 200 ms/100 pA. **(H)** Bar graphs of -Ih at the different ages under control and Rheb^CA^ conditions. Inset: Zoom of the control condition bar graphs. One way ANOVA. Data are mean ± SEM. ****:P<0.0001, ***:P<0.001, *:P<0.05, and ns: not significant. Exact *P* values can be found in Supplementary Table 4.

### Ectopic HCN4 expression in human TSC and FCDII neurons

Data presented above indicate that HCN4 channels confer a spiking advantage in Rheb^CA^ neurons that otherwise would not be able to generate repetitive firing upon cAMP stimulation. To examine whether abnormal HCN4 expression occurs in patients with FCM, we obtained cortical tissue from five patients that underwent surgery for epilepsy due to FCM. Three patients had the histopathological diagnosis of FCDII and two patients had TSC. Patients had FCM detected on MRI and underwent EEG with a combination of subdural grid and depth electrodes prior to FCM resection. Patients were identified as FCDII post-surgery based on pathological examination of the H&E-stained resected tissue and identification of hallmarks of FCDII, including cortical dyslamination and the presence of cytomegalic neurons. In addition, for tissue used for immunofluorescence, we confirmed the presence of cytomegalic neurons using immunostaining for SMI 311 (Fig. 6C, E, and Supplementary Fig. S5 for FCDII; Fig. 7D and Supplementary Fig. S6 for TSC), a pan-neuronal neurofilament (NF) that is accumulated in the soma and dendrites of dysmorphic neurons in human and murine FCD and is considered a marker of these neurons (12, 16, 42, 43). Dysmorphic SMI 311-immunopositive neurons expressed the neuronal markers, NeuN and neurofilament-light chain (NFL) (Supplementary Fig. S5). Consistent with previous reports, NFL was enriched in the soma of dysmorphic neurons that co-expressed SMI 311 (16). We also immunostained for phospho-S6 as a read-out of mTOR activity. SMI 311-immunopositive dysmorphic neurons displayed increased immunoreactivity for phospho-S6 compared to normal-appearing, surrounding cells, indicating mTOR hyperactivity (Supplementary Fig. S5 and S6). In all five tissue samples, we identified HCN4 immunoreactivity (green) in cytomegalic cells and no cellular staining in surrounding cells (Fig. 6 for FCDII and Fig. 7 for TSC). HCN4-positive fibers were visible in some samples and may correspond to either processes of diseased neurons or thalamic inputs or both (44). HCN4 immunostaining was performed with the two different antibodies also tested in mice as indicated on the figures. Cytomegalic cells expressing HCN4 displayed enlarged nuclei as shown by hematoxylin staining (in paraffin-embedded samples, Fig. 6A). In addition, cells displaying HCN4 immunoreactivity expressed increased phospho-S6 (Fig. 6B and D and Fig. A-C) and SMI 311 identifying them as cytomegalic, diseased neurons (Fig. 6C and E and Fig. 7D and E). Collectively, these data show that both mouse FCM neurons and dysmorphic human neurons from TSC and FCDII patients display abnormal expression of HCN4 channels.

**Figure 6:**
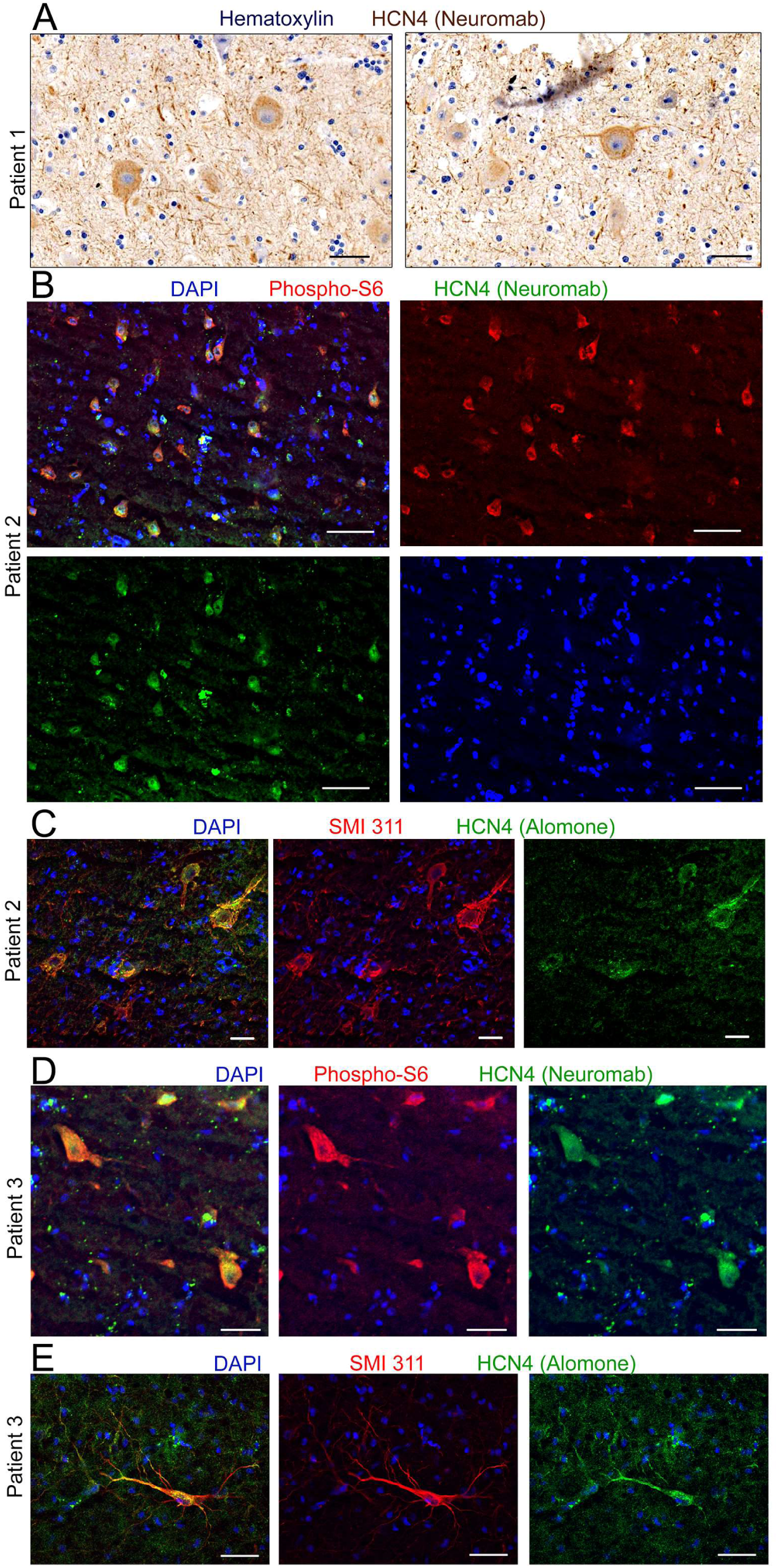
Ectopic HCN4 expression in diseased neurons in human FCDII cortices. **(A)** Staining for HCN4 in patient 1 with FCDII. Scale bar: 30 µm. **(B and C)** Immunostaining for HCN4 and phospho-S6 or SMI-311 in FCDII tissue from patient 2 co-stained with DAPI. Scale bars: 70 µm (B) and 30 µm (C). **(D and E)** Immunostaining for HCN4 and phospho-S6 or SMI-311 in FCDII tissue from patient 3 co-stained with DAPI. Scale bars: 35 µm (D) and 40 µm (E).

**Figure 7:**
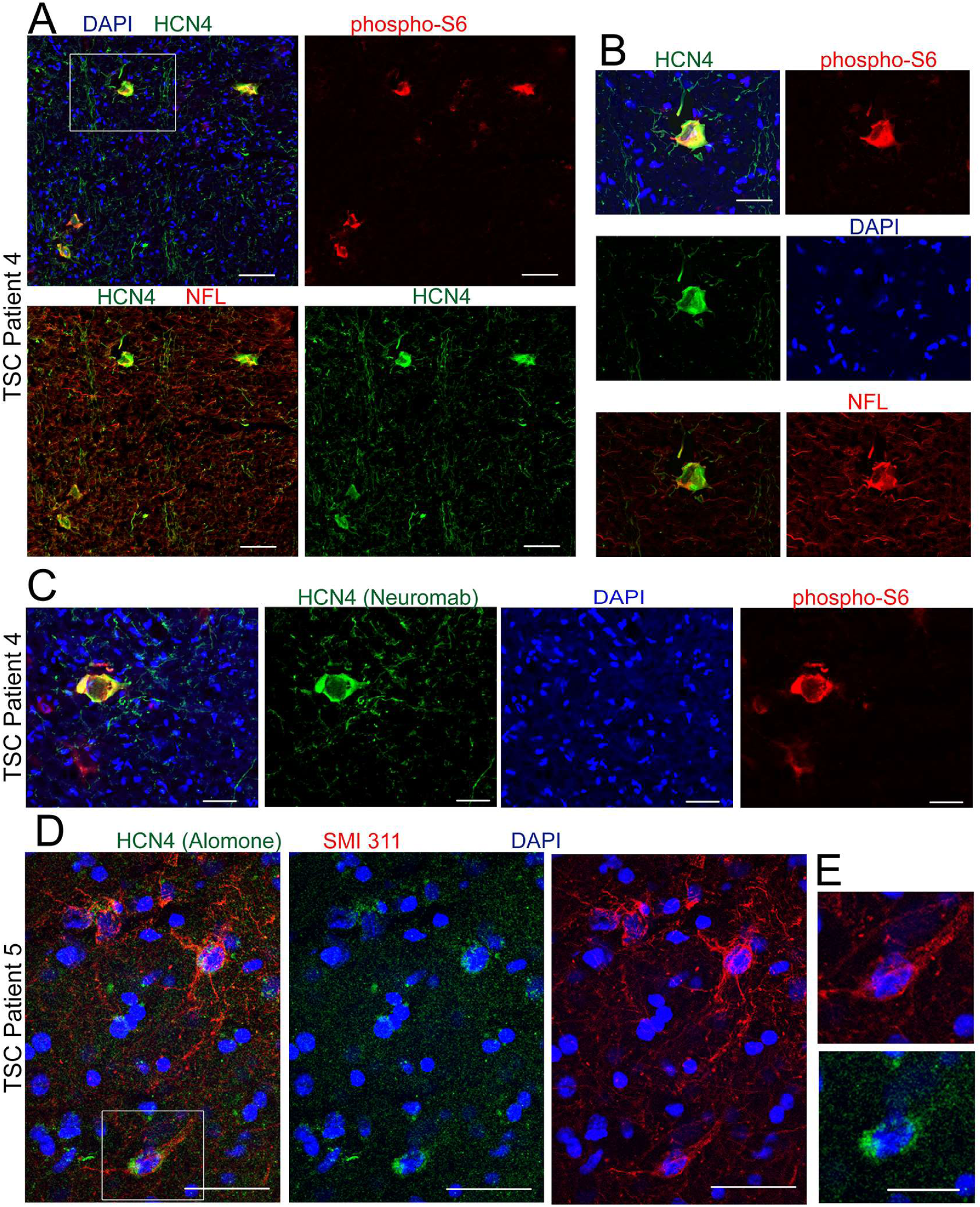
Ectopic HCN4 expression in diseased neurons in human TSC cortices. **(A and C)** Immunostaining for HCN4 and phospho-S6 TSC tissue from patient 4 co-stained with DAPI. Scale bars: 70 µm **(A)** and 40 µm **(C)**. **(B)** Higher magnification of the cell shown in the white square in A. Scale bar: 35 µm. **(D)** Immunostaining for HCN4 and SMI 311 in TSC tissue from patient 5 co-stained with DAPI. Scale bar: 40 µm. **(E)** Higher magnification of the cell shown in the white square in D. Scale bar: 20 µm.

### Blocking HCN4 activity prevents the establishment of epilepsy

HCN channels contribute to the generation of rhythmic firing in neurons and heart cells (31–34). In addition, data in Figure 3 showed that HCN4 channel expression drove firing in Rheb^CA^ neurons upon cAMP stimulation. To then directly address whether the ectopic expression of HCN4 channels in Rheb^CA^ neurons drives the generation of seizures, we designed a strategy to block HCN channel activity *in vivo*. We expressed nonfunctional HCN4 (HCN4^NF^) subunits that were generated by adding two mutations into HCN4 (45). HCN4^NF^ subunits are expected to form heteromers with endogenous HCN4 subunits and render the endogenous HCN4 channel unable to conduct ions. Indeed, both the sag responses observed in current clamp and the h currents recorded in voltage clamp in Rheb^CA^ neurons were eliminated in Rheb^CA^ neurons that co-expressed HCN4^NF^ (Fig. 8A and B). Furthermore, HCN4^NF^ expression significantly normalized the resting membrane potentials of Rheb^CA^ neurons (Fig. 8C) similarly to what was shown with the HCN channel blocker zatebradine in Figure 3. Similar to the experiments with Kir2.1, expressing HCN4^NF^ in Rheb^CA^ neurons did not interfere with mTOR hyperactivity as measured by phospho-S6 immunofluorescence and soma size (Fig. 8D and E). HCN4^NF^ expression in Rheb^CA^ neurons did not alter the properties of action potentials (Fig. 8F) or the input-output curve (Fig. 8G and H). Thus, expressing HCN4^NF^ in Rheb^CA^ neurons did not alter their ability to generate actions potentials, but it hyperpolarized them back to control levels and blocked the activity of endogenous HCN4 channels. Finally, a 7-day long continuous video-EEG monitoring of mice containing Rheb^CA^ neurons expressing HCN4^NF^ revealed that these mice had no seizures whereas littermate mice containing Rheb^CA^ neurons (without HCN4^NF^) displayed a mean of three daily, convulsive seizures (Fig. 8I and J). Together, these data indicate that the ectopic activation of HCN4 channels in FCM neurons are necessary for generating seizures.

**Figure 8:**
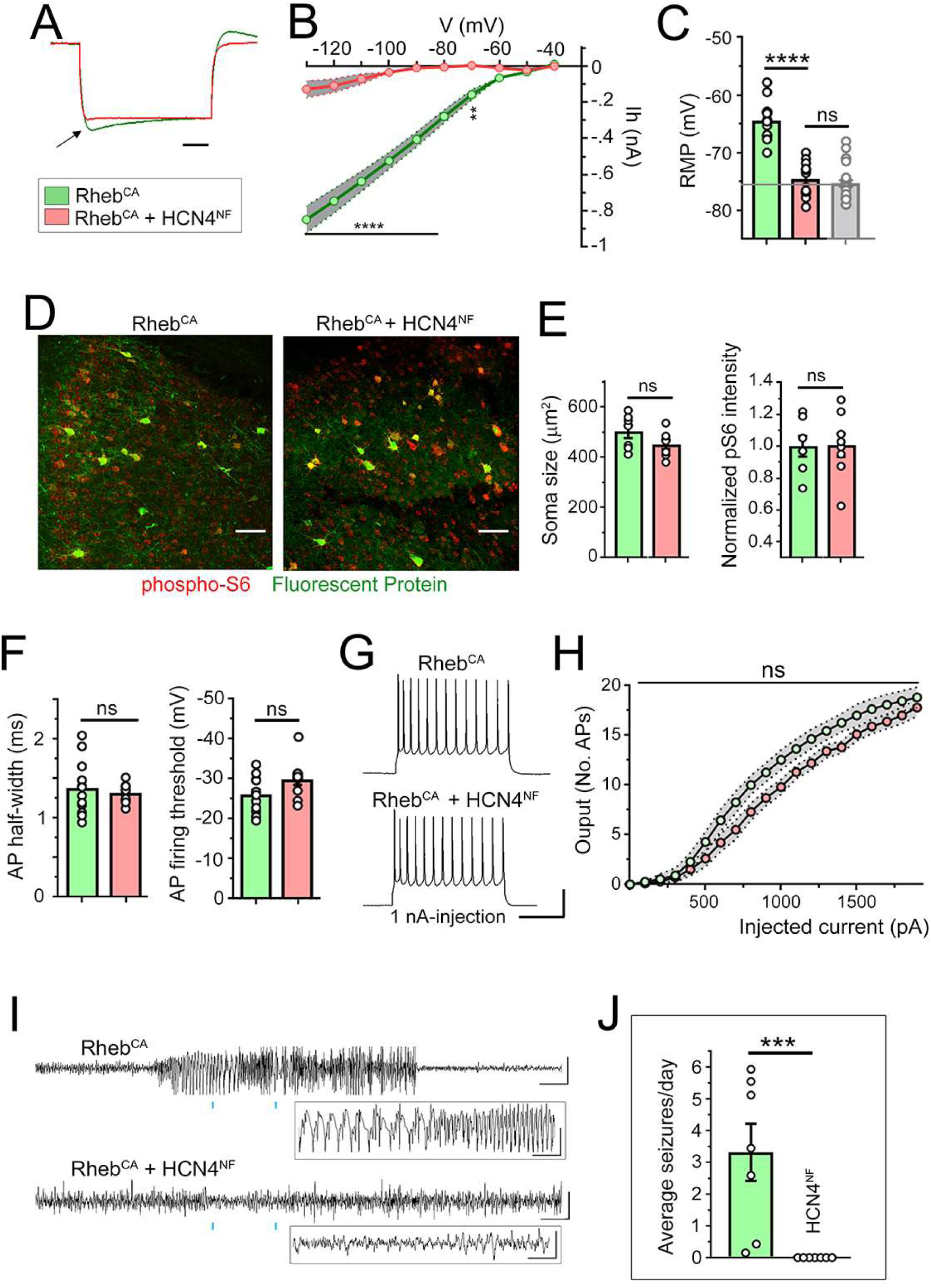
Blocking HCN4 channel activity in FCM neurons prevents epilepsy. **(A and B)** Representative voltage-traces of Rheb^CA^ neurons with and without nonfunctional HCN4 (HCN4^NF^) channels expression. Neurons were recorded in acute slices from P21-P35 littermate mice electroporated with Rheb^CA^+GFP or Rheb^CA^+HCN4^NF^ (+ tdTomato). (B) Ih-V curve for each condition. Two-way repeated measure ANOVA followed by Sidak post-test. **(C)** Bar graphs of RMP. The control from panel A in Fig. 2 (in grey) is added for comparison. Patch clamp recordings were obtained in acute slices from P21-P35 mice. Student’s t test. **(D)** Confocal images of GFP and tdTomato fluorescence (pseudo-colored green) and phospho-S6 immunostaining (red) in coronal sections from 4 months old mice expressing Rheb^CA^ + GFP or + HCN4^NF^ (+ tdTomato). Scale bars: 80 µm. **(E)** Bar graphs of soma size and normalized (to GFP control cells) phospho-S6 immunofluorescence for neurons expressing Rheb^CA^+GFP or Rheb^CA^+HCN4^NF^. Student’s t test. **(F)** Bar graphs of the action potential (AP) threshold and half-width. Student’s t test. **(G)** Representative depolarization and action potentials upon current injection in Rheb^CA^ neurons with and without HCN4^NF^. Scales: 200 ms/40 mV. **(H)** Input-output curves in Rheb^CA^ neurons with and without HCN4^NF^. Two-way repeated measure ANOVA followed by Sidak post-test. The grey area indicates the SEM for each curve. **(I)** Representative EEG traces in Rheb^CA^ mice with and without HCN4^NF^. **(J)** Bar graphs of the number of seizures per day. Mann Whitney U test. Data are mean ± SEM. *: P<0.05, **: P<0.01, ***: P<0.001, ****: P<0.0001, ns: not significant. Exact *P* values can be found in Supplementary Table 4.

## Discussion

In light of previous data reporting depolarized RMPs of Rheb^CA^ neurons, we hypothesized that Rheb^CA^ neurons may be more excitable and thus trigger seizures. To test this hypothesis, we expressed Kir2.1 channels in Rheb^CA^ neurons to silence them and examine whether this would limit seizure activity. Kir2.1 expression hyperpolarized the RMPs of Rheb^CA^ neurons and increased their membrane conductance leading to a right shift in the input-output curve without altering their increased cell size, mTOR hyperactivity, and misplacement. These data show that Rheb^CA^ neurons expressing Kir2.1 were indeed less excitable by being hyperpolarized and less responsive to depolarization. As a result of this manipulation, mice had significantly fewer seizures per day, suggesting that Rheb^CA^ neurons are more excitable than control neurons and trigger seizures. We next obtained whole cell patch clamp recordings of Rheb^CA^ neurons to further investigate the mechanism responsible for their increased excitability. We confirmed that Rheb^CA^ neurons have depolarized RMPs and are thus closer to the threshold for generating action potentials. However, they also displayed increased conductance and thus required a larger depolarizing current injection to reach firing threshold, hence Rheb^CA^ neurons were less excitable in response to depolarization than control neurons. These findings initially seemed to contradict the conclusion from the Kir2.1 experiment that Rheb^CA^ neurons could trigger seizures. However, we further identified the aberrant expression of HCN currents that provided an explanation for the unusual, increased likelihood of Rheb^CA^ neurons to generate action potentials independent of depolarizing inputs as detailed below. In young adults, Rheb^CA^ neurons expressed larger inward currents than in controls, including a combination of HCN- and presumed Kir-mediated currents. Outward currents could not be examined in Rheb^CA^ neurons due to poor space clamp leading to unclamped firing upon depolarizing voltage pulses. Due to the large increase in Rheb^CA^ neuron size, it was not surprising to find larger Kir-mediated currents, which are well-known to be expressed in control layer 2/3 pyramidal neurons as G protein inwardly rectifying K channels (46, 47). However, finding HCN-mediated currents was surprising because control layer 2/3 pyramidal neurons are not known to express such currents, as confirmed here. HCN channels, in particular HCN2 and HCN4, are sensitive to cAMP levels with higher cAMP levels leading to increased HCN-mediated currents (17–19). Considering that the levels of cAMP increases in developing hippocampal neurons (doubled from P6 to P15) (48) and could be altered in disease condition, it is conceivable that the increase in HCN currents could also be due to increased cAMP levels in Rheb^CA^ neurons compared to control neurons. We could not directly measure cAMP levels in Rheb^CA^ neurons. However, using immunostaining we identified the selective HCN4 expression that was not present in control neurons andexpressing HCN4^NF^ led to a significant reduction of HCN-mediated current. These data suggest that HCN-mediated currents in Rheb^CA^ neurons are primarily due to the ectopic expression of HCN4 channels independent of changes in cAMP levels. With respect to neuron excitability, we found that increasing intracellular cAMP levels with forskolin was sufficient to trigger the firing of Rheb^CA^ neurons. Therefore, although Rheb^CA^ neurons have an increased membrane conductance and require more current injection to reach firing threshold, they are more depolarized and an increase in intracellular cAMP levels, as opposed to excitatory input-induced depolarization, acts as the trigger to induce Rheb^CA^ neuron firing. These data show an unanticipated and novel mechanism of excitability that is consistent with a significant decrease in the excitatory drive (i.e., frequency of spontaneous excitatory synaptic inputs) onto Rheb^CA^ neurons (21). Identifying a cAMP-dependent excitability in Rheb^CA^ neurons is tantalizing because cortical pyramidal neurons receive multiple inputs that activate receptors leading to cAMP increases. These inputs include noradrenergic and dopaminergic innervation from the locus coeruleus (e.g., (49, 50)) and the ventral tegmental area (e.g., Gs-coupled β1- and β2-adrenergic receptors and D1 and D5 receptors, see (47, 51) for references). Noradrenaline or dopamine released from these inputs would thus contribute to cAMP increases and HCN4 channel activation that could lead to firing and seizures It is also conceivable that inhibitors of these receptors, preventing increases in intracellular cAMP levels, may alleviate seizure activity.

Regarding the identity of the HCN channels expressed in Rheb^CA^ neurons, we found the selective expression of HCN4 channels by immunostaining. It is possible that Rheb^CA^ neurons also acquire HCN1 or HCN2 channels at densities below detection levels for immunostaining. Nevertheless, the important finding was the fact that HCN4 expression was absent in control pyramidal neurons in our sections and has not been reported in cortical pyramidal neurons in young adult mice despite the presence of mRNA neurons (35–40). We found that about 85% of the Rheb^CA^ neurons labeled with tdTomato displayed HCN4 immunoreactivity. Using co-electroporation of two plasmids, we routinely achieved colocalization of the two plasmids in 92% of the electroporated cells (52). As such we underestimate the percent of cells expressing HCN4 by 8% leading to an adjusted total of about 93% of the cells expressing HCN4. Considering that every recorded Rheb^CA^ neuron expressed HCN-mediated currents, albeit with different amplitude, it is possible that HCN4 expression in a subset of Rheb^CA^ neurons was below the detection limit for immunostaining. Finding HCN4 channels in neurons that normally do not express these channels may seem surprising. However, as shown in the developing hippocampus (48, 53), HCN4 is strongly expressed perinatally in the cortex and is dramatically decreased in young adults (P30) (37). HCN4 channels are thus intrinsic to cortical neurons during development. Considering HCN4 expression was mTOR-dependent, and mTOR increases protein translation, it is conceivable that the ectopic HCN4 expression results from increased translation of mRNA already present in pyramidal cortical neurons.

Our findings add to a large body of literature on HCN expression and seizures. Indeed, several studies have reported alterations in the expression of HCN, in particular, HCN1 and HCN2, in different types of epilepsy in both mice and humans (for review (41, 54)). One recent study reported that blocking HCN1 channel activity prevented absence seizures (55), suggesting that an increase in HCN1 channel expression contributes to the generation of absence seizures. Another study reported no change in HCN4 expression in the hippocampus following febrile seizures despite changes in HCN1 (56). Loss-of-function mutations in HCN4 has also recently been associated with benign myoclonic epilepsy (57). More recently, increased HCN4 expression in the hippocampal dentate gyrus has been reported in individuals with SUDEP (58). However, it was unclear whether the alterations in increased HCN4 expression preceded seizure occurrence and were responsible for seizures. Our model of FCD-associated seizures allowed us to demonstrate that HCN4 expression is uniquely expressed in dysmorphic neurons and precedes seizures. Indeed, patch clamp recordings reported the abnormal expression of HCN currents in Rheb^CA^ neurons as early as P8, about two weeks prior to the onset of convulsive seizures (observed starting at P21). In addition, no HCN4 expression was detected in the contralateral hemispheres lacking FCM, but experiencing epileptiform activity. Finally, we show clear causality between HCN4 activity and seizure occurrence, as genetically blocking HCN4 activity prevented the development of seizures.

Identifying ectopic HCN4 has several important clinical implications. At the present time, there are no HCN blockers that are selective for HCN4. However, our findings support HCN4 as a prime candidate for shRNA-based gene therapy for treating seizures associated with FCDII and TSC that exhibit focal cortical malformations. While ectopic HCN4 expression was eliminated with rapamycin treatment, directly targeting HCN4 for epilepsy treatment would prevent the severe adverse events that occur when using the higher rapamycin doses necessary to improve efficacy (7). Furthermore, considering that HCN4 is downstream of mTOR signaling, our findings are likely applicable to other mTORopathies resulting from mutations in the mTOR and GATOR pathway genes (e.g., AKT, PI3K, mTOR, RHEB, DEPDC5, NRPL2/3 (3, 4, 59–71).

In conclusion, we have provided evidence that enlarged, dysmorphic mutant neurons in mouse and human FCMs express HCN4 channels that are normally absent in cortical neurons in young adults. This ectopic HCN4 channel expression is mTOR-dependent, precedes the development of epilepsy, and contributes to the generation of seizures by enhancing the excitability of FCM neurons. Our findings add to the body of literature on HCN channels in epilepsy, and more importantly, highlight a novel mechanism of seizures by the unexpected contribution of the isoform 4 of HCN channels that has high cAMP-sensitivity. This mechanism can explain how sensory stimulations leading to the activation of specific cAMP-generating dopaminergic or adrenergic inputs onto FCM neurons would trigger seizures. In addition, the unique expression of HCN4 channels in dysmorphic FCM neurons provides a highly specific target for gene therapy for epilepsy treatment for individuals with TSC and FCDII.

## Materials and Methods

### Animals

Research protocols were approved by the Yale University Institutional Animal Care and Use Committee. All experiments were performed on CD-1 (Charles River), an outbred strain of mice of either sex.

### *In utero* electroporation and plasmids

Each DNA plasmid (Supplementary Table 2) was diluted in sterile PBS (pH 7.4) to a final concentration of 1.5-3 μg/μl (specific concentrations below). About 1 μl of DNA solution containing 0.1% fast green was injected into the lateral ventricle of E15.5±0.5 fetuses with a glass pipette. After injection, PBS soaked tweezer-type electrodes (model 520, BTX) were positioned on heads of the fetuses across the uterine wall and 6 square-pulses (42V, 50 ms duration, 950 ms intervals) were applied using a pulse generator (ECM830, BTX). Mice were prescreened for successful electroporation on a fluorescence enabled stereo microscope (SZX16, Olympus) prior to recruitment for EEG monitoring. To generate epilepsy-associated FCM, a DNA solution composed of pCAG-Rheb^S16H^ (Rheb^CA^, 2 μg/μl) and pCAG-tdTomato (1 μg/μl) was injected in half of all fetuses while pCAG-GFP (3 μg/μl) containing solution was injected in the remaining littermates as controls. To silence FCM neurons, pCAG-Kir2.1-T2A-tdTomato (2 μg/μl) + pCAG-Rheb^CA^ (2 μg/μl) was injected in half of the fetuses while the remaining half received pCAG-Rheb^CA^ (2 μg/μl) + pCAG-GFP (2 μg/μl) as controls.

### Seizure detection and analysis

Animals were randomly assigned an arbitrary identification number without knowing the experimental condition before implanting dual independent channel EEG electrodes (Fig. 1B) for identification after the double-blind analyses. Six-pin EEG headmounts (made inhouse) were attached with two stainless steel machine screws and a dab of cyanoacrylate to the skulls of electroporated animals of >2 months of age. After one week of recovery, EEG preamplifiers were attached to the implants and to an electrical commutator (Pinnacle Technology Inc.) to allow tethered recordings from freely moving animals. EEG (sampled at 400 Hz) were recorded with digital video continuously for 7 consecutive days for each animal. Epileptiform activity was analyzed post hoc, while blinded to the condition, using Sirenia Seizure Pro software (Pinnacle Technology Inc.) to identify possible seizure epochs. An automated line length search method was applied to all recorded EEG channels with the threshold set at 5000 length/sec using a 10 second search window with a 0.5 second sliding window. Identified episodes were verified manually with video inspection. Convulsions that reached Racine stage 3-5 (72), from forelimb clonus to rearing and falling with forelimb clonus, were counted as a seizure. Seizure duration was defined from the onset of convulsion to cessation of all motor movement. Data are reported as average number of seizures per day. Power spectrums, with and without temporal component, were analyzed for each epoch, from the onset of convulsion to the cessation of motor movements. For preictal (baseline) power spectrum analyses, with and without temporal component, corresponding time window 60 seconds before convulsion onset was used (Pinnacle Seizure Pro software).

### Brain slice preparation and Immunohistochemistry

Mice were deeply anesthetized with pentobarbital (50 mg/kg) and perfused transcardially with ice cold phosphate buffered saline (PBS, pH 7.4) followed by ice cold paraformaldehyde (PFA, 4%). Perfused brains were also drop fixed in 4% PFA for an additional hour after removal from the skull. Fixed brains were cryoprotected with 30% sucrose in PBS overnight at 4°C and serially sectioned into 50 μm thick sections using a freezing microtome. Sections were blocked for 1 hour at room temperature in blocking buffer consisting of 2% BSA and 0.3% Triton X-100 in PBS. Floating sections were incubated overnight at 4°C in primary antibodies diluted in blocking buffer. Following 3 washes in PBS and an additional 15 min in blocking solution at room temperature, sections were then incubated with secondary antibodies at 1:1000 dilution for 2 hours at room temperature. ProLong Gold antifade reagent (Life Technologies) was used to mount and preserve stained sections. While blinded to the experimental conditions, Z-stack images were acquired using a fluorescence confocal microscope (FV1000, Olympus) with a 20x dry objective (N.A. 0.75, Olympus) and reconstructed using Imaris 4.0 (Bitplane AG) and Photoshop CS6. Fluorescence intensities and cell sizes were quantified using ImageJ 1.39t (Freeware, Wayne Rasband, NIH). Cell size was quantified by averaging 15 brightest fluorescently tagged cells per animal. Immunofluorescence of phospho-S6 in the same 15 brightest cells was quantified using integrated fluorescence. Antibodies were previously validated by previous studies.

The list of antibodies is detailed in Supplementary Table 3. Antibodies were extensively used and validated by previous studies. In addition, the specificity of the HCN4 antibody was further confirmed by RNAi in vitro (Supplementary Fig. S2) and exclusive immunoreactivity in neurons electroporated with an HCN4 overexpression plasmid and not in non-electroporated neurons in the vicinity (Supplementary Fig. S3).

### Acute slice preparation and whole cell recording

1-5-week-old mice were used for acute slices. Tissue was dissected and sliced in ice cold artificial cerebral spinal fluid (aCSF) oxygenated with 95% O_2_/5%CO_2_. The aCSF contained (all in mM): 124 NaCl, 3 KCl, 1.25 NaH_2_PO_4_, 1 MgSO_4_, 26 NaHCO_3_, 10 Dextrose, 2 CaCl_2_, 0.4 ascorbate, 4 Na-Lactate, 2 Na-Pyruvate (290±5 mOsm/kg, pH 7.2). Coronal sections (350 µm) were prepared using a vibratome (Vibratome 1000). Sections were incubated in aCSF at 32°C for 45 minutes before returning to room temperature (25°C) where they were kept for 8 to 10 hours during experimentation. Fluorescent neurons in the mPFC were visualized using epifluorescence on an Olympus BX51WI microscope with a 40X water immersion objective (Olympus, LUMPlanFL/IR). Whole-cell recordings were performed at 28°C using pulled glass pipettes (4-7 ΩM) filled with internal solution (in mM: 110 K-gluconate, 4 KCl, 10 HEPES, 10 di-tris-phosphocreatine, 4 Mg-ATP, 0.3 Na-GTP). Recordings were acquired with an amplifier (Axopatch 200B, Molecular Devices). Measured values were not adjusted for the liquid-junction potential. The resting membrane potential was recorded within the first 10 seconds after achieving whole-cell configuration in current-clamp mode, while the cell is at rest without any holding current. The membrane conductance was calculated using the average membrane potential change from ten hyperpolarizing current injections of −500 pA in current-clamp mode when the cell is at rest. The action potential half width was calculated at half of the peak amplitude of a single action potential induced by current injection. The action potential firing threshold was defined as the membrane potential at which the first derivative of an action potential in current-clamp mode achieves 10% of its peak velocity (dV/dt). The action potential input-output curve was generated by injecting positive current in current-clamp mode from 0 to 1.4 or 2 nA at 50 or 100 pA increments for 500 msec.

### Human tissue sample

Human tissue was obtained from different sources: directly from the surgery room or from the Tuberous sclerosis Alliance. Freshly resected tissue was rapidly frozen in 2-Methylbutane (isopentane; Acros Organics MS; AC12647-0010) at <-80°C, rapidly frozen in liquid nitrogen and then stored at −80°C (for immunoblotting and immunostaining), or fixed in 10% formalin (for immunohistochemistry). The sample from the TSA alliance was provided as frozen. Human tissue collection and use was approved by the human ethical committee in Xiangya Hospital, Central South University, China and by the Yale University human ethical committee. Freshly frozen tissue was sectioned (15 µm) via a cryostat and fixed in 4% PFA for 2 minutes before immunostaining. Patient information is provided in Supplementary Table 1.

### IMR-32 (human neuroblastoma) cell line and immunoblotting

IMR-32 cells were grown at 37°C with 5% CO_2_ in Eagle’s Minimum Essential Medium supplemented with 10% fetal bovine serum and 1% penicillin-streptomycin. Cells were plated in 6-well culture plates and transfected when they reached ∼70% confluence. Transfection of the HCN4 overexpression plasmid was done using Lipofectamine 3000 reagent (Invitrogen) and transfection of the Stealth RNAi siRNAs against HCN4 [Invitrogen: HSS1125216, HSS115218, HSS173331 (human) MSS221131 (mouse)] was done using Lipofectamine RNAiMAX reagent (Invitrogen) according to the manufacturer’s protocol. Cells were collected 48 hours after transfection for western blotting. Sample lysates were resolved with SDS-PAGE, transferred onto PVDF, blocked in 5% milk, and incubated with primary antibodies (concentrations of primary antibody are listed in Supplementary Table S3) overnight. HRP-conjugated anti-rabbit or anti-mouse were used as secondary antibodies.

### Statistical analyses

All analyses were conducted blindly knowing only the arbitrarily assigned animal ID (independent of electroporation condition). Statistical tests and plots were performed using Prism 7 (GraphPad Software, Inc.), with P < 0.05 for significance for all experiments. Specific statistical tests used for each experiment are described in Supplementary Table 4. Data are presented as mean ± SEM.

## Supplementary Materials

Fig. S1: The concentration of Rheb^CA^ influences h current amplitudes.

Fig. S2: RNAi validation of HCN4 antibodies.

Fig. S3: HCN4 antibodies detect HCN4 overexpression.

Fig. S4: HCN4 immunostaining in the cerebellum.

Fig. S5: Immunostaining in human FCDII tissue

Fig. S6: Immunostaining in human TSC tissue

Table S1: Patient information.

Table S2: Constructs used.

Table S3: Primary and Secondary antibodies.

Table S4: Summary of statistical tests.

Movie S1: Example of a convulsive seizure in a mouse containing a FCM

## Acknowledgments

We thank Drs. Stieber and Ludwig (Institute of Experimental and Clinical Pharmacology and Toxicology, Erlangen, Germany) for the pcDNA3-hHCN4 plasmid and Drs. Hanada and Maehama (National Institute of Infectious Diseases) for the Rheb^CA^ plasmid.

## Funding

This work was supported by NIH/NINDS R21 NS093510, DoD TS150058, R01 NS111980, and the Swebilius Foundation (AB), Brown Coxe Postdoctoral Fellowship and American Epilepsy Society Postdoctoral Fellowship, and NIH/NICHD F32 HD095567 (LHN), and National Natural Science Foundation of China 81671123 (LZ). We thank the Tuberous Sclerosis Alliance to provide one TSC sample.

## Author contributions

LH and AB designed the experiments. LSH, JW, LHN, and LZ performed the experiments. DS, JTR, and LZ provided human tissue samples. AB, LH and LHN wrote the manuscript. Competing interests: We have no conflict of interest.

