## Supplementary Material for "Ectopic HCN4 expression drives mTOR-dependent epilepsy"

**Supplementary Materials:**

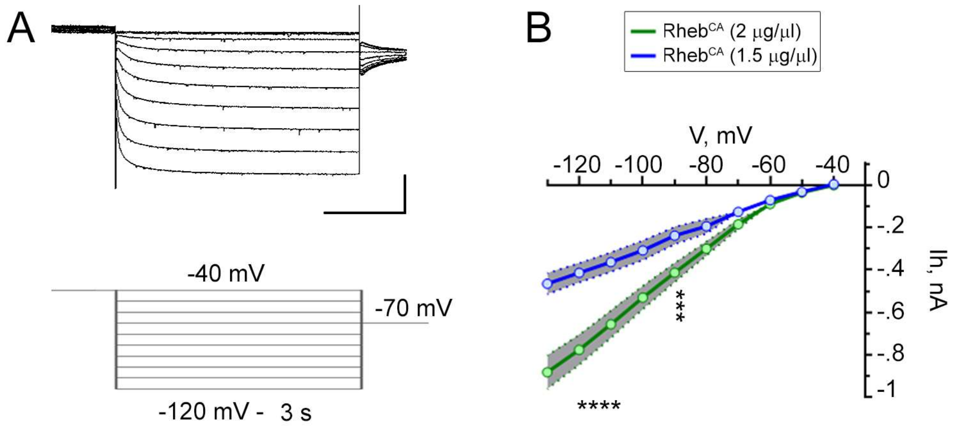


**Supplementary Fig. 1: The concentration of Rheb$^{CA}$ influences h current amplitudes. (A)** 3 s-long voltage pulse to fully activate HCN currents in Rheb$^{CA}$ neurons. **(B)** Current-voltage (Ih-V) curve obtained in P35-P40 neurons containing Rheb$^{CA}$ at 1.5 µg/µl (blue) or at 2 µg/µl (green, same as in Fig. 3E). Two-way repeated measures ANOVA followed by Sidak post-test, **** P<0.0001 and ***:P<0.001. n=8 and 20 neurons for Rheb$^{CA}$ 1.5 and 2.0 µg/µl, respectively. The grey areas illustrate the SEM for each curve. The exact statistical parameters are listed in Supplementary Table 4.

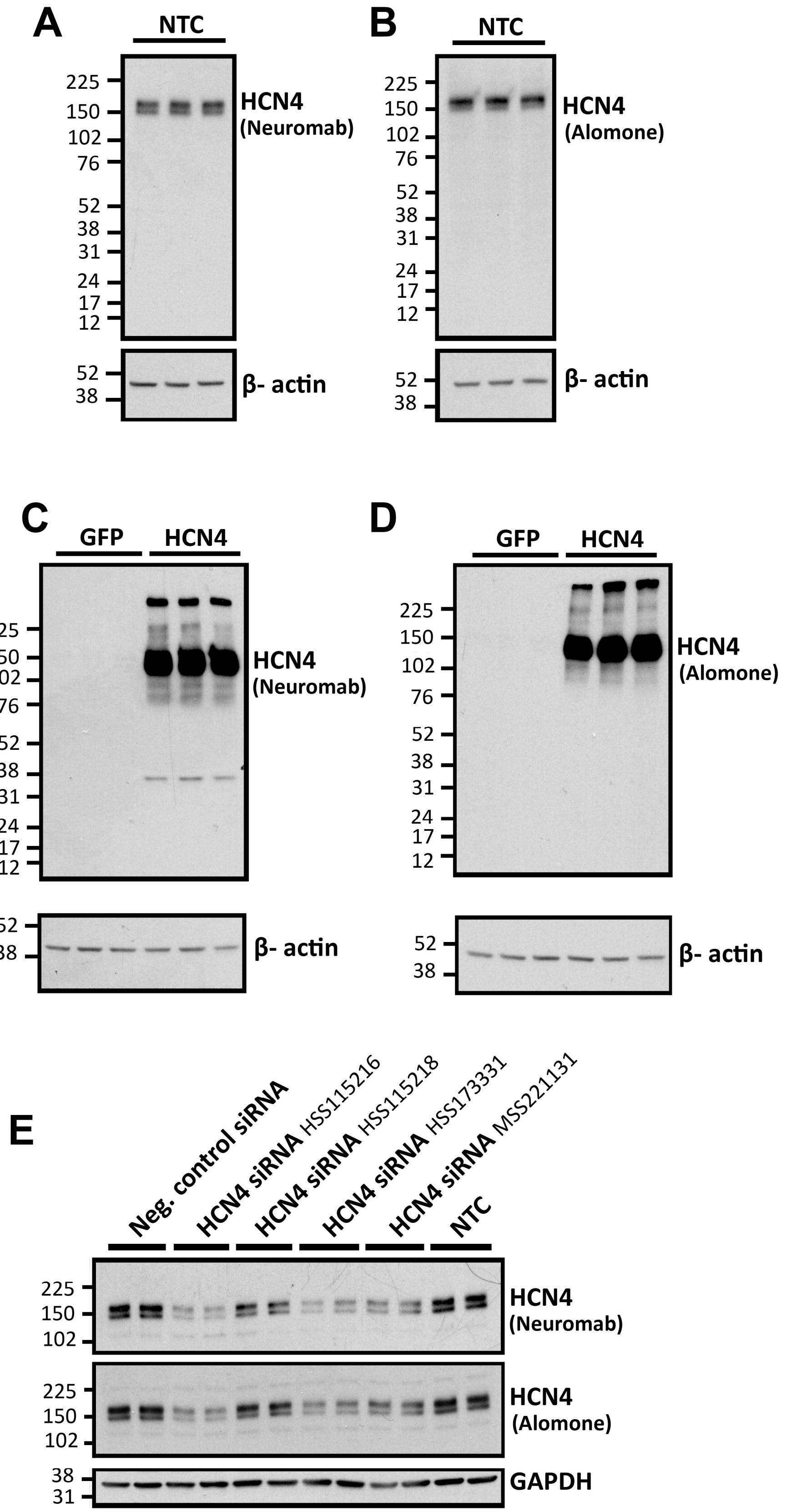
A
NTC
HCN4
(Neuromab)
β- actin
B
NTC
HCN4
(Alomone)
β- actin
C
GFP
HCN4
HCN4
(Neuromab)
β- actin
D
GFP
HCN4
HCN4
(Alomone)
β- actin
E
Neg. control siRNA
HCN4 siRNA HSS115216
HCN4 siRNA HSS115218
HCN4 siRNA HSS173331
HCN4 siRNA MSS221131
NTC
HCN4
(Neuromab)
HCN4
(Alomone)
GAPDH

**Supplementary Fig. 2: Validation of HCN4 antibody specificity in IMR-32 (human neuroblastoma) cell line. (A-E)** Western blots of HCN4 from lysates from non-transfected cells (A and B), lysates from cells following HCN4 overexpression (C and D), and lysates from cells following HCN4 knockdown (Invitrogen Stealth RNAi siRNAs) (E). Blots were probed with either a mouse HCN4 antibody (1:1000, Neuromab #75-150) (A, C, E) or a rabbit HCN4 antibody (1:500-1:1000, Alomone #APC-052) (B, D, E). β-actin (1:2000, Cell Signaling #4970) and GAPDH (1:5000, Cell Signaling #5174) were used as loading controls. Experiments were ran in triplicates (A-D) and duplicates (E). *NTC, non-transfected cells.*

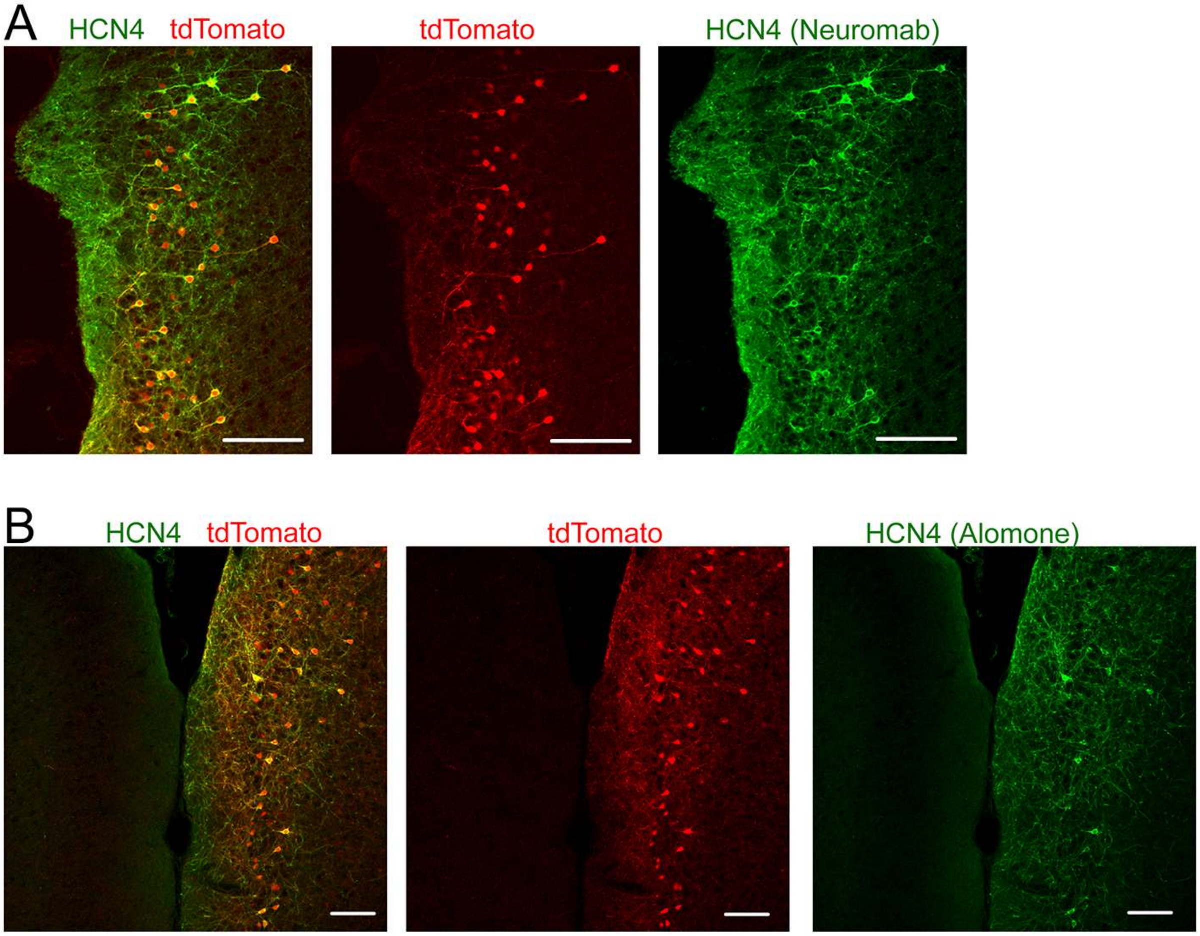


**Supplementary Fig. 3: HCN4 antibody detects HCN4 overexpression**. **(A and B**) HCN4 immunostaining (with Neuromab (A) and Alomone (B) antibodies) and tdTomato fluorescence in coronal sections from 2 months old mice containing cortical neurons electroporated with pCAG-tdTomato and pCAG-HCN4 at E15. Scale bars: 100 µm (A) and 70 µm (B).

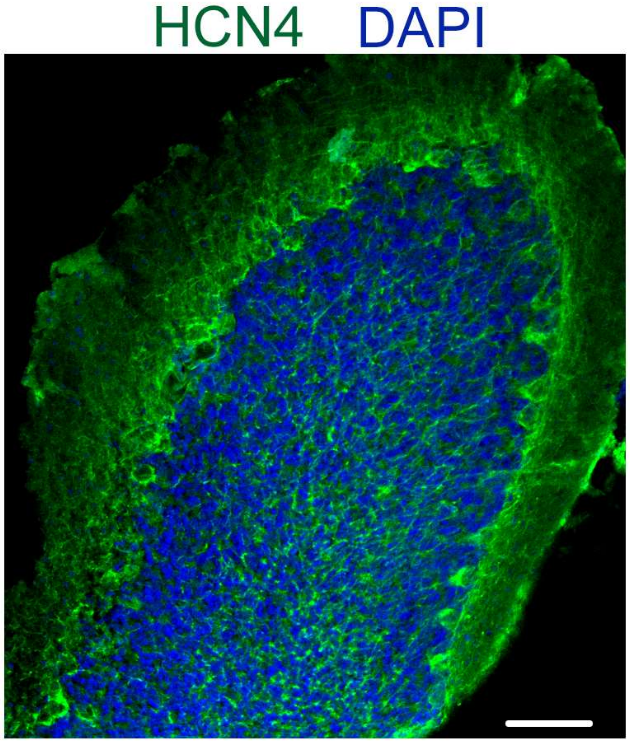


**Supplementary Fig. 4: HCN4 immunostaining in the cerebellum.** HCN4 (Neuromab antibody) and DAPI staining in cerebellar staining of 2 months old mice.

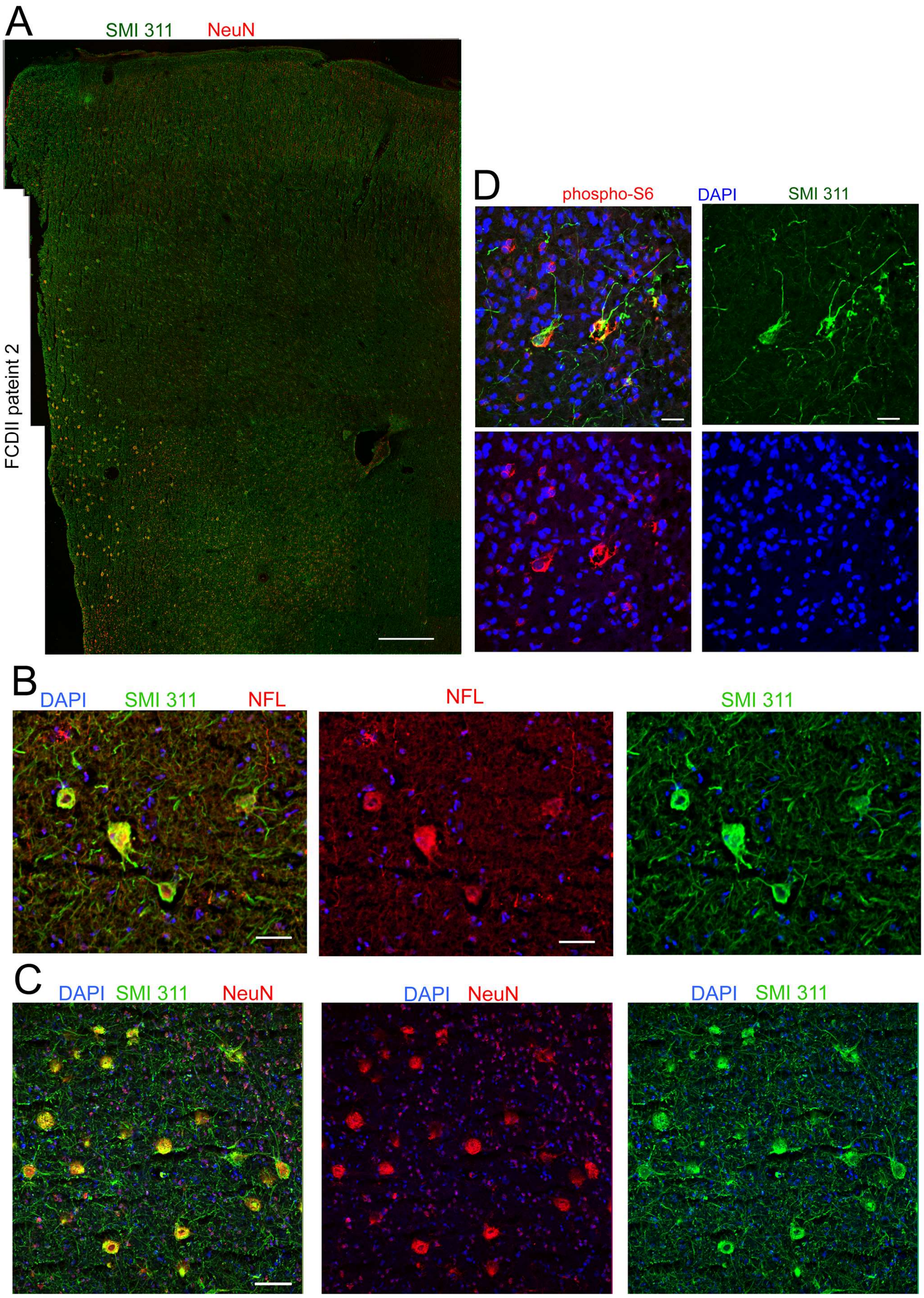
A
SMI 311
NeuN
FCDII pateint 2
D
phospho-S6
DAPI
SMI 311
B
DAPI
SMI 311
NFL
NFL
SMI 311
C
DAPI
SMI 311
NeuN
DAPI
NeuN
DAPI
SMI 311

**Supplementary Fig. 5: Identification of cytomegalic neurons in FCDII. (A)** A confocal micrograph montage of cortical tissue stained with SMI 311 and NeuN from a patient with FCDII (patient 2). Scale bar: 630 µm. **(B and C)** Immunostaining for SMI311 and either NeuN (B) or NFL (C) in FCDII tissue (patient 2). DAPI is a nuclear stain. Scale bars: 150 µm (NeuN) and 40 µm (NFL). **(D)** Immunostaining for SMI 311 and phospho-S6 in FCDII tissue co-stained with DAPI from patient 3. Scale bar: 30 µm

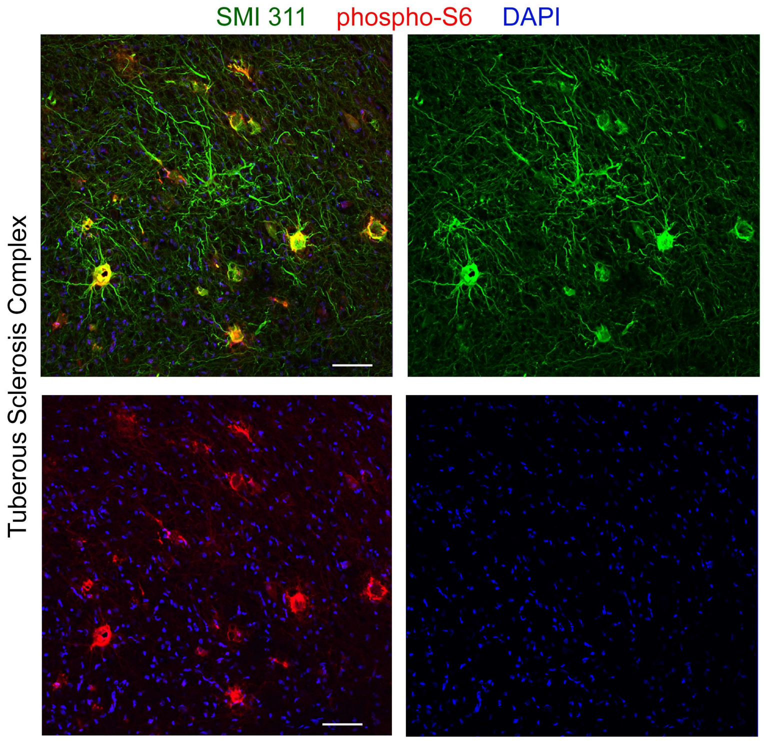


**Supplementary Fig. 6: Identification of cytomegalic neurons in TSC.** Immunostaining for SMI 311 and phospho-S6 in TSC tissue co-stained with DAPI. Scale bar: 70 µm.

**Supplementary Table 1: Patient information**

| Patient # | Age | Sex | Type | Medical notes |
|---|---|---|---|---|
| 1 | 5 | Male | FCD IIb | Left superior frontal gyrus seizures, cortical dyslamination, dysmorphic neurons, balloon cells, and gliosis |
| 2 | 35 | Female | FCD IIa | Intractable Focal Impaired Awareness (FIA) seizures from right occipital temporal lobes, cortical dyslamination, dysmorphic neurons |
| 3 | 12 | Male | FCD IIa | Left frontal focal motor seizures intractable, cortical dyslamination, dysmorphic neurons |
| 4 | 4 | Female | TSC | Right frontal opercular tuber |
| 5 | 21 | Female | TSC | Focal seizures, cortical tuber and subependymal nodules |

**Supplementary Table 2: Constructs used.**

| Plasmid | IUE concentration (µg/µL) | Notes/Origin |
|---|---|---|
| pCAGGS-Rheb S16H (Rheb$^{CA}$) | 2.0; 1.5 (for Fig. 1) | Gift from T. Maehama and K. Hanada (National Institute of Infectious Diseases, Tokyo, Japan) (*1*) |
| pCAG-GFP | 1.0 – 4.0 (as noted in figure legend) | Addgene (catalog no.11150) (*2*) |
| pCAG-tdTomato | 1.0 | Addgene (#83029) (*3*) |
| pCAG-Kir2.1-T2A-tdTomato | 2.0 | Addgene (#60598) (*4*) |
| pCAGEN-hHCN4-AYA (HCN4$^{NF}$) | 4.0 | Synthesized coding sequence for hHCN4 (AJ132429.1) with G480A/G482A mutations (*5*), inserted into EcoRI and XhoI sites of pCAGEN (Addgene; catalog no. 11160). |
| pCAGEN-hHCN4 | 2.0 | Subcloned hHCN4 from pcDNA3-hHCN4 (*6*) into EcoRI and XhoI sites of pCAGEN (Addgene; catalog no. 11160). |
| pcDNA3-hHCN4 | N/A | Gift from J. Stieber and A. Ludwig (Institute of Experimental and Clinical Pharmacology and Toxicology, Erlangen, Germany) (*6*) |
| pCAGEN | N/A | Addgene (catalog no. 11160) |

**Supplementary Table 3: Primary and Secondary antibodies**

| Antibody | Company | Catalog Number | Host animal | Concentration used in IHC or WB when specified |
|---|---|---|---|---|
| Primary | | | | |
| HCN1 | UC Davis/NIH NeuroMab Facility | N70/28 (75-110) | Mouse | 1:750 |
| HCN2 | UC Davis/NIH NeuroMab Facility | N71/37 (75-111) | Mouse | 1:750 |
| HCN3 | UC Davis/NIH NeuroMab Facility | N141/28 (75-175) | Mouse | 1:750 |
| HCN4 | UC Davis/NIH NeuroMab Facility | N114/10 (75-150) | Mouse | 1:750<br>1:250 (human)<br>1:1000 WB |
| HCN4 | Alomone Labs | APC-052 | Rabbit | 1:300-1:500<br>1:500-1:1000 WB |
| NeuN | EMD Millipore | ABN78 | Rabbit | 1:2000 |
| NeuN | EMD Millipore | MAB 377 | Mouse | 1:2000 |
| NF-L | Cell Signaling | C28E10 | Rabbit | 1:500 |
| NF-L | Abcam | ab134460 | Chicken | 1:1000 |
| pS6 (S240/244) | Cell Signaling | D68F8 | Rabbit | 1:4000 |
| SMI311 | Covance | SMI-311R | Mouse | 1:4000 |
| SMI32 (NfH) | Abcam | 8135 | Rabbit | 1:750 |
| β-actin | Cell Signaling | 4970 | Rabbit | 1:2000 WB |
| GAPDH | Cell Signaling | 5174 | Rabbit | 1:5000 WB |
| DAPI | Life Technologies | D1306 | | 1:36000 |
| Secondary | | | | |
| α Mouse IgG Alexa Fluor 633 | Thermo Fisher Scientific | A-21052 | Goat | 1:1000 |
| α Mouse IgG Alexa Fluor 488 | Thermo Fisher Scientific | A-11001 | Goat | 1:1000 |
| α Mouse IgG Alexa Fluor 568 | Thermo Fisher Scientific | A-11004 | Goat | 1:1000 |
| α Mouse IgG Alexa Fluor 555 | Thermo Fisher Scientific | A28180 | Goat | 1:1000 |
| α Rabbit IgG Alexa Fluor 488 | Thermo Fisher Scientific | A-11034 | Goat | 1:1000 |
| α Rabbit IgG Alexa Fluor 555 | Thermo Fisher Scientific | A-31572 | Donkey | 1:1000 |
| α Rabbit IgG Alexa Fluor 647 | Thermo Fisher Scientific | A-31573 | Donkey | 1:1000 |
| α Chicken IgY Alexa Fluor 488 | Thermo Fisher Scientific | A-11039 | Goat | 1:1000 |
| α Chicken IgG Alexa Fluor 555 | Thermo Fisher Scientific | A-21437 | Goat | 1:1000 |

| | | | | |
|---|---|---|---|---|
| α Chicken IgG Alexa Fluor 647 | Thermo Fisher Scientific | A-21449 | Goat | 1:1000 |

WB: western blot

### Supplementary Table 4: Summary of statistical tests

| Figure | Test | Statistical value | P value | n | Definition of n |
|---|---|---|---|---|---|
| ***Figure 1*** | | | | | |
| 1F, soma size | Student's t test, unpaired, two-tailed | t (19)=23.14 | <0.0001 | 10 (GFP)<br>11 ($Rheb^{CA}$) | Number of mice |
| 1G, phospho-S6 | Student's t test, unpaired, two-tailed | t (6)=5.994 | <0.001 | 4 (GFP)<br>4 ($Rheb^{CA}$) | Number of mice |
| ***Figure 2*** | | | | | |
| 2C, pS6 intensity | Student's t test, unpaired, two-tailed | t (11)=1.118 | 0.287 | 6 ($Rheb^{CA}$)<br>7 ($Rheb^{CA}$ + Kir2.1) | Number of mice |
| 2C, cell size | Student's t test, unpaired, two-tailed | t (11)=0.2957 | 0.773 | 6 ($Rheb^{CA}$)<br>7 ($Rheb^{CA}$ + Kir2.1) | Number of mice |
| 2D, Capacitance | Student's t test, unpaired, two-tailed | t (18) =0.8511 | 0.406 | 10 ($Rheb^{CA}$)<br>10 ($Rheb^{CA}$ + Kir2.1) | Neurons from 3 mice ($Rheb^{CA}$) and 3 mice ($Rheb^{CA}$ + Kir2.1) |
| 2D, RMP | Student's t test, unpaired, two-tailed | t (18)=3.0820 | 0.0064 | 10 ($Rheb^{CA}$)<br>10 ($Rheb^{CA}$ + Kir2.1) | Neurons from 3 mice ($Rheb^{CA}$) and 3 mice ($Rheb^{CA}$ + Kir2.1) |
| 2D, Conductance | Student's t test, unpaired, two-tailed | t (21)=3.448 | 0.0024 | 12 ($Rheb^{CA}$)<br>11 ($Rheb^{CA}$ + Kir2.1) | Neurons from 5 mice ($Rheb^{CA}$) and 2 mice ($Rheb^{CA}$ + Kir2.1) |
| 2F, AP half-width | Student's t test, unpaired, two-tailed | t (25)=0.00714 | 0.9944 | 12 ($Rheb^{CA}$)<br>15 ($Rheb^{CA}$ + Kir2.1) | Neurons from 5 mice ($Rheb^{CA}$) and 3 mice ($Rheb^{CA}$ + Kir2.1) |
| 2F, AP firing threshold | Student's t test, unpaired, two-tailed | t (23)=0.5570 | 0.5825 | 12 ($Rheb^{CA}$)<br>13 ($Rheb^{CA}$ + Kir2.1) | Neurons from 5 mice ($Rheb^{CA}$) and 3 mice ($Rheb^{CA}$ + Kir2.1) |
| 2H | Two-way repeated measures ANOVA Sidak post-test | Interaction:<br>F (29, 580)=3.364, p<0.0001<br>Row factor:<br>F (29, 580)=47.08, p<0.0001<br>Column factor:<br>F (1, 20)=6.279, p=0.0219<br>Subjects (matching)<br>F (20, 580)=18.16, p<0.0001 | 0.05 | 8 ($Rheb^{CA}$)<br>14 ($Rheb^{CA}$ + Kir2.1) | Neurons from 2 mice ($Rheb^{CA}$) and 3 mice ($Rheb^{CA}$ + Kir2.1) |
| 2J, seizure duration | Student's t test, unpaired, two-tailed | t (8)=0.292 | 0.7777 | 7 ($Rheb^{CA}$)<br>3 ($Rheb^{CA}$ + Kir2.1) | Number of mice |
| 2J, number of seizures per day | Mann Whitney U test, two-tailed | U=3 | 0.0082 | 7 ($Rheb^{CA}$)<br>6 ($Rheb^{CA}$ + Kir2.1) | Number of mice |
| ***Figure 3*** | | | | | |
| 3A, RMP | Student's t test, unpaired, two-tailed | t (45)=12.15 | <0.0001 | 26 (GFP)<br>21 ($Rheb^{CA}$) | Neurons from 5 mice (GFP) and 4 mice ($Rheb^{CA}$) |
| 3B, conductance | Student's t test, unpaired, two-tailed | t (24)=13.46 | <0.0001 | 15 (GFP)<br>11 ($Rheb^{CA}$) | Neurons from 3 mice (GFP) and 3 mice ($Rheb^{CA}$) |

| | | | | | |
|---|---|---|---|---|---|
| 3D | Two-way repeated measures ANOVA Sidak post-test | From 0 to 500 pA injection:<br>Interaction:<br>F (5, 90) = 14.57, p<0.0001<br>Row factor:<br>F (5, 90) = 90.44, p<0.0001<br>Column factor:<br>F (1, 18) = 14.51, p=0.0013<br>Subjects (matching)<br>F (18, 90) = 11.51, p<0.0001<br>From 600 to 1200<br>F (6, 66) = 0.5321, P=0.78<br>F (6, 66) = 129.7, P<0.0001<br>F (1, 11) = 18.05, P=0.0014<br>F (11, 66) = 57.67, P<0.0001 | 0.05 | From 0 to 500 pA injection<br>11 (GFP)<br>9 ($Rheb^{CA}$)<br><br>From 600 to 1200 pA injection<br>4 (GFP)<br>9 ($Rheb^{CA}$) | Neurons from 4 mice each condition |
| 3H | Two-way repeated measures ANOVA Tukey post-test | Interaction:<br>F (18, 90) = 40.75, p<0.0001<br>Row effect:<br>F (9, 90) = 229.4, p<0.0001<br>Column effect:<br>F (2, 10) = 26.61, p<0.0001<br>Subjects (matching):<br>F (10, 90) = 9.292, p<0.0001 | 0.05 | 5 (GFP)<br>5 ($Rheb^{CA}$)<br>3 ($Rheb^{CA}$ + Zatebradine) | Neurons from 2 mice (GFP),<br>4 mice ($Rheb^{CA}$),<br>and<br>2 mice ($Rheb^{CA}$ + Zatebradine) |
| 3I, RMP ($Rheb^{CA}$ vs $Rheb^{CA}$ +Zatebradine) | Student's t test, unpaired, two-tailed | t (16)=8.294 | <0.0001 | 11 ($Rheb^{CA}$)<br>7 ($Rheb^{CA}$+ Zatebradine) | Neurons from 3 mice (GFP) and 3 mice ($Rheb^{CA}$) |
| 3J | Two-way repeated measures ANOVA Tukey post-test | Interaction:<br>F (18, 288)=27.03, p<0.0001<br>Row effect:<br>F (9, 288)=44.72, p<0.0001<br>Column effect:<br>F (2, 32)=31.71, p<0.0001<br>Subjects (matching):<br>F (32, 288)=14.43, p<0.0001 | 0.05 | 8 (GFP)<br>20 ($Rheb^{CA}$)<br>7 ($Rheb^{CA}$ + Zatebradine) | Neurons from 2 mice (GFP),<br>4 mice ($Rheb^{CA}$),<br>and<br>2 mice ($Rheb^{CA}$ + Zatebradine) |
| 3K | Pearson correlation | r=-0.6887 | 0.0008 | 20 | Neurons from 7 mice |
| 3M | Student's t test, paired, two-tailed | t (8)=4.16 | 0.0032 | 9 (ACSF)<br>9 (Forskolin in ACSF)<br>8 (ACSF)<br>8 (Forskolin in ACSF) | $Rheb^{CA}$ neurons from 3 mice<br><br>GFP neurons from 3 mice |
| ***Figure 5*** | | | | | |
| 5F | Two-way repeated measures ANOVA Tukey post-test | Interaction:<br>F (9, 180)=35.26, p<0.0001<br>Row effect:<br>F (9, 180)=59.49, p<0.0001<br>Column effect:<br>F (1, 20)=39.64, p<0.0001<br>Subjects (matching):<br>F (20, 180)=10.79, p<0.0001 | 0.05 | 9 (GFP)<br>13 ($Rheb^{CA}$) | Neurons from 2 mice (GFP),<br>3 mice ($Rheb^{CA}$), |
| *5H* | One-way ANOVA Tukey post-hoc | F (3,77)=64.23<br>P<0.0001 | 0.042, GFPvsRheb$^{CA}$ <P12<br>>0.99, GFP, <P12 vs>P28<br><0.0001, Rheb$^{CA,}$ <P12vs>P28 | 24 (<P12 GFP)<br>14 (>P28 GFP)<br>25 (<P12 $Rheb^{CA}$)<br>20 (>P28 $Rheb^{CA}$) | Each group from >3 mice |
| ***Figure 8*** | | | | | |

| | | | | | |
|---|---|---|---|---|---|
| 5B | Two-way repeated measures ANOVA Sidak post-test | Interaction: F (9,153)=62, p<0.0001 Row factor: F (9, 153)=106, p<0.0001 Column factor: F (1, 17)=100.1, p=0.0001 Subjects (matching) F (17, 153)=9.188, p<0.0001 | 0.05 | 8 ($Rheb^{CA}$) 11 ($Rheb^{CA}$ + $HCN4^{NF}$) | Neurons from 4 mice ($Rheb^{CA}$) 3 mice ($Rheb^{CA}$ + $HCN4^{NF}$) |
| 5G, RMP | Student's t test, unpaired, two-tailed | t (21)=7.412 | <0.0001 | 12 ($Rheb^{CA}$) 11 ($Rheb^{CA}$ + $HCN4^{NF}$) 26 (control from Fig 2) | Neurons from 4 mice ($Rheb^{CA}$) and 3 mice ($Rheb^{CA}$ + $HCN4^{NF}$) |
| 5E, pS6 intensity | Student's t test, unpaired, two-tailed | t (13)=0.08209 | 0.9358 | 7 ($Rheb^{CA}$) 8 ($Rheb^{CA}$ + $HCN4^{NF}$) | Number of mice |
| 5E, soma size | Student's t test, unpaired, two-tailed | t (12)=1.534 | 0.1510 | 7 ($Rheb^{CA}$) 7 ($Rheb^{CA}$ + $HCN4^{NF}$) | Number of mice |
| 5F, AP half-width | Student's t test, unpaired, two-tailed | t (2)=0.5903 | 0.5616 | 12 ($Rheb^{CA}$) 10 ($Rheb^{CA}$ + $HCN4^{NF}$) | Neurons from 4 mice ($Rheb^{CA}$) and 3 mice ($Rheb^{CA}$ + $HCN4^{NF}$) |
| 5F, AP firing threshold | Student's t test, unpaired, two-tailed | t (20)=2.011 | 0.0580 | 12 ($Rheb^{CA}$) 10 ($Rheb^{CA}$ + $HCN4^{NF}$) | Neurons from 4 mice ($Rheb^{CA}$) and 3 mice ($Rheb^{CA}$ + $HCN4^{NF}$) |
| 5H | Two-way repeated measures ANOVA Sidak post-test | Interaction: F (29, 551)=2.276, p<0.0002 Row factor: F (29, 580)=371.2, p<0.0001 Column factor: F (1, 19)=2.905, p=0.1046 Subjects (matching) F (19, 551)=23.65, p<0.0001 | 0.05 | 11 ($Rheb^{CA}$) 10 ($Rheb^{CA}$ + $HCN4^{NF}$) | Neurons from 3 mice ($Rheb^{CA}$) and 3 mice ($Rheb^{CA}$ + $HCN4^{NF}$) |
| 5J | Mann Whitney U test | U=0 | 0.0006 | 7 ($Rheb^{CA}$) 7 ($Rheb^{CA}$ + $HCN4^{NF}$) | Number of mice |
| ***Supplementary Figures*** | | | | | |
| S1 | Two-way repeated measures ANOVA Sidak post-test | Interaction: F (9, 234)=8.988, p<0.0001 Row factor: F (9, 234)=88.68, p<0.0001 Column factor: F (1, 26)=7.072, p=0.0132 Subjects (matching) F (26, 234)=14.99, p<0.0001 | 0.05 | 8 ($Rheb^{CA}$, 1.5 µg/µl 20 ($Rheb^{CA}$, 2 µg/µl) | Neurons from 2 mice ($Rheb^{CA}$, 1.5 µg/µl) and 4 mice ($Rheb^{CA}$, 2 µg/µl, used in Fig. 2e) |
